# Quantitative evidence for foraging preferences of shrill carder bumble bee (*Bombus sylvarum*) queens

**DOI:** 10.64898/2026.10.21.760793

**Authors:** Julia de Korab Skulska, Corneile Minnaar, Mark JF Brown

## Abstract

**Introduction:** Bumblebees are facing declines globally due to agricultural intensification and climate change. In the UK, the shrill-carder bumble bee, *Bombus sylvarum*, has dramatically declined and is now restricted to five isolated populations across England and Wales. A key life-cycle stage for bumblebees is the spring queen, who is responsible for colony establishment, but we lack quantitative data to inform conservation practice for this life stage.

**Aim/methods:** To quantify foraging preferences and floral resource dependency of spring queens in *B. sylvarum.* We determined the floral resources preferred by *B. sylvarum* queens across their activity period, along with the potential for inter-specific resource competition.

**Results:** At the start of the season, *B. sylvarum* queens foraged solely on *Lamium album* and *Trifolium repens*. Later they increased their foraging breadth but predominantly foraged on *Vicia villosa* and *Rhinanthus minor*. Plant–pollinator networks indicate the potential for competition with more common bumble bee species, with *B. sylvarum* queens exhibiting a broader foraging range than these potential competitors.

**Discussion:** As the season progresses, the foraging diversity of *B. sylvarum* queens increases. Later in the season, the lower species specificity index for *B. sylvarum* queens suggests that these queens are less specialised than their congeneric competitors.

**Implications for insect conservation:** Our results show that early in the season *B. sylvarum* queens rely on two floral species. Managing sites to maintain and increase strong populations of these plant species will likely enhance queen success, which is crucial to enable colony establishment and population growth.

**Statements and Declarations:** The authors have no competing interests. This research was funded as part of the MSc by Research degree at Royal Holloway, University of London. This research was funded, in part, by a UKRI grant from the Natural Environment Research Council (NERC) NE/Y000285/1.

## Introduction

Insects are facing declines across the globe caused by climate change, pollution, and land use change (Wagner 2020; Wagner et al. 2021; Neff et al. 2022). In turn, this puts the ecosystem services that insects provide (Losey and Vaughan 2006) at risk. For example, 75% of the world’s food crop cultivars (Klein et al. 2007), and 87.5% of flowering plant species (Ollerton, Winfree, Tarrant 2011) are dependent on animal pollination, which is predominantly provided by insects, and in particular, bees. Consequently, local and global declines in bee populations are a significant concern (Buchmann and Ascher 2005; Winfree et al. 2009; Potts et al. 2010; Nieto et al. 2014; Powney et al. 2019; Dicks et al. 2021; Zattara and Aizen 2021).

Bumblebees, a major group of insect pollinators, are not immune to these declines (Williams 1986; Williams and Osborne 2009; Cameron and Sadd 2020; Soroye et al. 2020). However, not all bumblebees are equally threatened, with threat status often linked to particular traits. For example, 8 of the 24 UK bumblebee species have long tongues, and 6 of these are threatened, whereas the more widespread and abundant species typically have short tongues (Bumblebee Conservation Trust 2026). Leguminous plants, which often bear flowers with deep corollas, provide a vital resource for long-tongued bumblebees. The loss of leguminous plants from crop rotations because of agricultural intensification may therefore be especially detrimental to long-tongued bumblebees (Williams 1986; Goulson et al. 2005; Goulson et al. 2008; Williams and Osborne 2009). Whether this is the major driver of declines in long-tongued bumblebees across landscapes is debated (Goulson et al. 2005; Williams 2005), and, likely, both climate specialisation and habitat specialisation (e.g., food requirements) are relevant to understanding why some species decline more than others (Williams 2005).

One long-tongued bumblebee facing decline in the UK is *Bombus sylvarum* (Williams 1986; Page et al. 2020; Horsley 2024). Although *B. sylvarum* was once widespread in the UK (Sladen 1912; Williams 1986), it is now the rarest bumblebee species in England and Wales, found in only five isolated populations across the south of the UK: Somerset, Pembrokeshire, the Gwent Levels, Port Talbot to Kenfig, and the Thames Estuary (Horsley 2024). These areas are primarily found near coasts, where agriculture is less practised and less intensive (Ellis et al. 2006). *B. sylvarum* favours florally rich habitats comprised of unimproved grasslands (Carvell 2002; Goulson et al. 2005; Benton 2006). Unimproved grasslands in the UK have decreased in area by over 90% between 1932 and 1984, a significant loss of habitat that may be driving *B. sylvarum* declines (Fuller 1987; Carvell et al. 2006).

The plants that *B. sylvarum* forages on are also facing declines in the UK, e.g., the leguminous Fabaceae family (Carvell et al. 2006). Fabaceae and Scrophulariaceae can comprise up to 76% of *B. sylvarum* visits to flowers within a community (Connop 2007), reflecting their importance to this endangered bumblebee species. However, while the plant species and families that *B. sylvarum* forages on are broadly known, we lack quantitative data on their foraging activity and preferences. This is particularly true for the earliest stage of the *B. sylvarum* lifecycle, spring queens, the success of which is essential for colony establishment. *B. sylvarum* queens forage on fewer plant families than workers and males (Connop 2007; Benton 2006) and may be more vulnerable to decline than other bumblebees because of their late emergence time of late April to May (Benton 2006; Horsley 2024). By the time *B. sylvarum* queens emerge, other bumblebee species have already established colonies with workers, leaving *B. sylvarum* queens to compete with entire colonies for resources (Williams 1989). It is therefore crucial for *B. sylvarum* queens to have access to floral resources that they can exploit with less interspecific competition (i.e., flowers with deep corollas). Furthermore, climate change may be driving earlier queen emergence from hibernation (Horsley 2024), leading to a mismatch with key floral resources (Goulson et al. 2005). Consequently, a quantitative understanding of the foraging requirements of *B. sylvarum* spring queens is crucial to developing management strategies for this essential stage of the species’ lifecycle.

In this study, we quantified the foraging activity and preferences of *B. sylvarum* queens at Coalhouse Fort, in south Essex. This site is located along the Thames Estuary, where one of the five remaining *B. sylvarum* populations in the UK is located (Horsley 2024). This study aimed to understand foraging preferences and resource competition in *B. sylvarum* queens to inform floral resource management. We investigated queen foraging preferences by mapping floral abundances and queen foraging visits. Queens were then followed in patches of preferred floral resources to understand the dynamics of foraging within patches and how this varies with patch size. We recorded floral visits and generated a plant–pollinator network to determine whether *B. sylvarum* queens might face competition for resources. Finally, we ran null model simulations on a subset of these plant–pollinator networks focused on *Bombus* species only, to test for floral-abundance-corrected preference for (and relative avoidance of) different plant species among bumblebees.

## Materials and methods

### Study area

*Bombus sylvarum* is a priority conservation species restricted to five populations in the UK, with the largest population located along the Thames estuary in England (across Essex and Kent) (Fig. 1a, b) (Horsley 2024). In February 2025, we explored six sites in Essex that are known to support *B. sylvarum* populations: Hadleigh Castle Country Park, Canvey Wick, Coalhouse Fort, Wat Tyler Country Park, Rainham Marshes, and Two Tree Island (Connop 2007).

**Fig. 1.**
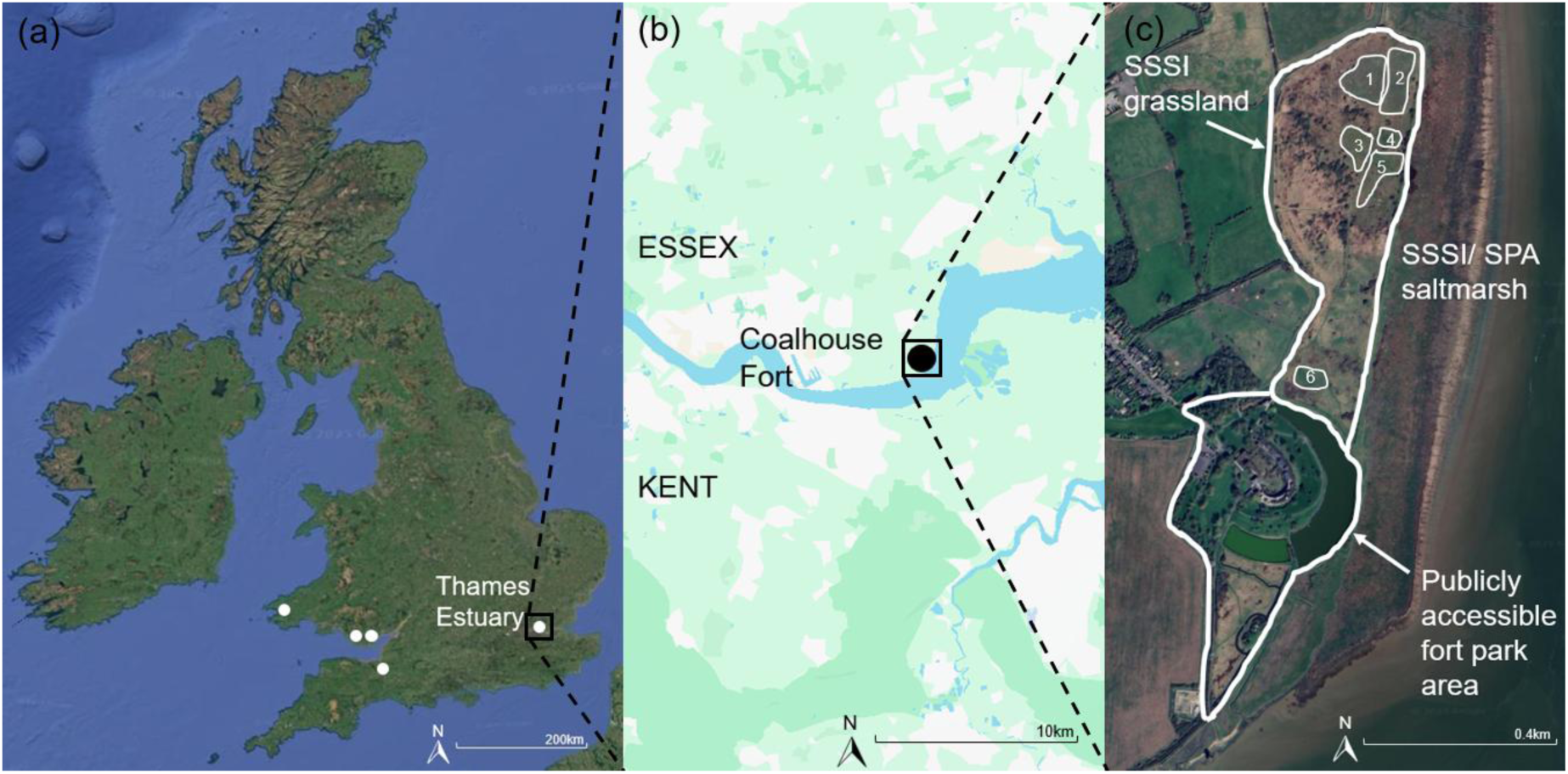
**(a)** The location of the five remaining *B. sylvarum* populations in the UK (from left to right): Pembrokeshire (Wales), Port Talbot to Kenfig (Wales), Gwent Levels (Wales), Somerset (England), and the Thames Estuary (England) (Horsley 2024). **(b)** the Thames Estuary site consists of Essex and Kent; the location of Coalhouse Fort is indicated as the black circle in Essex. **(c)** the publicly accessible parts of Coalhouse Fort consist of a park area around the fort, and a grassland that is considered a Site of Special Scientific Interest (SSSI); the SSSI grassland at Coalhouse Fort contains six meadows (see numbered polygons) (Google Earth 2025)

Coalhouse Fort (51°27’52.29”N, 0°26’1.05”W) was chosen for our study because it had suitable floral resources and a good count of *B. sylvarum* queens in 2024 (32 queens) (Reeves and Headley, pers. comm., 2024) (Fig. 1b). The other sites explored were either too large to survey effectively, or had low levels of flowering plants, so were not feasible for surveys during the spring. Coalhouse Fort consists of a publicly accessible park area directly surrounding the fort, and a grassland classed as a Site of Special Scientific Interest (SSSI, incorporating rank grassland, lichen heath, and calcareous grassland), formerly known as the East Tilbury Silt Lagoons (Fig. 1c) (Thurrock Council 2013). While this grassland is also open to the public, access is restricted to designated footpaths.

Data collection was divided into two phases. In the first phase, we collected *B. sylvarum* queen floral preference and abundance data at the publicly accessible park area around Coalhouse Fort. In the second phase, we generated plant–pollinator networks along transects within the meadows in the SSSI grassland.

### Data collection in the publicly accessible park area around Coalhouse Fort

Data collection took place in May 2025, once *B. sylvarum* queens were observed by Ray Reeves, the ranger at Coalhouse Fort. Shrill carder queen surveys were carried out between 09:00-18:00 GMT, when temperatures were above 15°C if it was cloudy, and above 13°C if it was sunny (following guidelines for UK Pollinator Monitoring Scheme (PoMS) FIT counts (UK Pollinator Monitoring Scheme 2023).

### Quantifying *B. sylvarum* queen floral preferences

The plants around the fort were surveyed before the meadows (Fig. 1c) because they flowered earlier in the year. We identified flowering plants using the Pl@ntNet application (Version 3.22.2b, Apple phone platform) and mapped the nine flowering species. To quantify insect visitation, we observed 1m² quadrats in monofloral patches of each plant species for 5-minute periods and recorded all visiting insects. Bumble bees were identified to species level, and where possible, solitary bees were identified to genus level using Falk and Lewington’s “Field guide to bees” (2015). We identified butterflies to species level using the “Wild ID guide for Butterflies of Britain and Ireland” (Bebbington 2024). Other pollinators were not identified to species level, but grouped by similar morphology, (e.g., ‘small black flies’). Observations were made for all nine plant species (see below for exceptions) within each hour of the 09:00-18:00 observation period. Observations took place across five days. The same patches were visited for each plant species within a day, but across days, different patches for each flowering plant species were randomly selected for observations. Flowering plant species patches were visited in a different order within the hourly slots each day, to account for temporal variation of flower-visiting insects. The only exceptions to these general rules were *Tragopogon porrifolius* and *Trifolium repens*. *T. porrifolius* flowers shut after noon, so observations were only possible in the mornings. As *T. repens* started flowering after the initial mapping of plant species, we took advantage of the afternoon slots for *T. porrifolius* observations to observe insect visitation to *T. repens* instead. In addition, there was only one patch of *T. porrifolius* at the site, so all observations of this species occurred at this patch, but different parts of the patch were selected on each day for the observations.

### Quantifying floral abundance of *Lamium album* and *Trifolium repens*

*Bombus sylvarum* queens at Coalhouse Fort were only observed to forage on *T. repens* and *L. album* (see Results). To quantify the availability of these resources, we identified all patches of each of these flowering species around the park area of Coalhouse Fort. We found twenty-seven *L. album* patches, and five *T. repens* patches (Fig. 2). Patch areas were estimated by calculating the area of the shape each patch was most similar too (e.g., rectangle or oval). Within each of the *T. repens* patches, we measured total floral abundance by counting individual flower heads as single floral units. We estimated the number of floral units per *L. album* patch using the patch area, the number of plants in a 1m² quadrat, and the average number of flowers per plant. Ten random plants were selected in each *L. album* patch and their flowers counted. We then calculated the average number of flowers per plant and counted the number of plants in a 1 m² quadrat of the patch. This was then scaled up to the total area of the patch to calculate the total floral abundance in each patch. *L. album* floral units were counted at the start of each day of 5-minute observations, except the first day (missed due to human error). *T. repens* floral units were calculated after the 5-minute observations, by counting flower heads in photos taken at the start of each day. *L. album* and *T. repens* floral units were calculated again, before 10-minute observations, to account for any temporal changes in abundance.

**Fig. 2.**
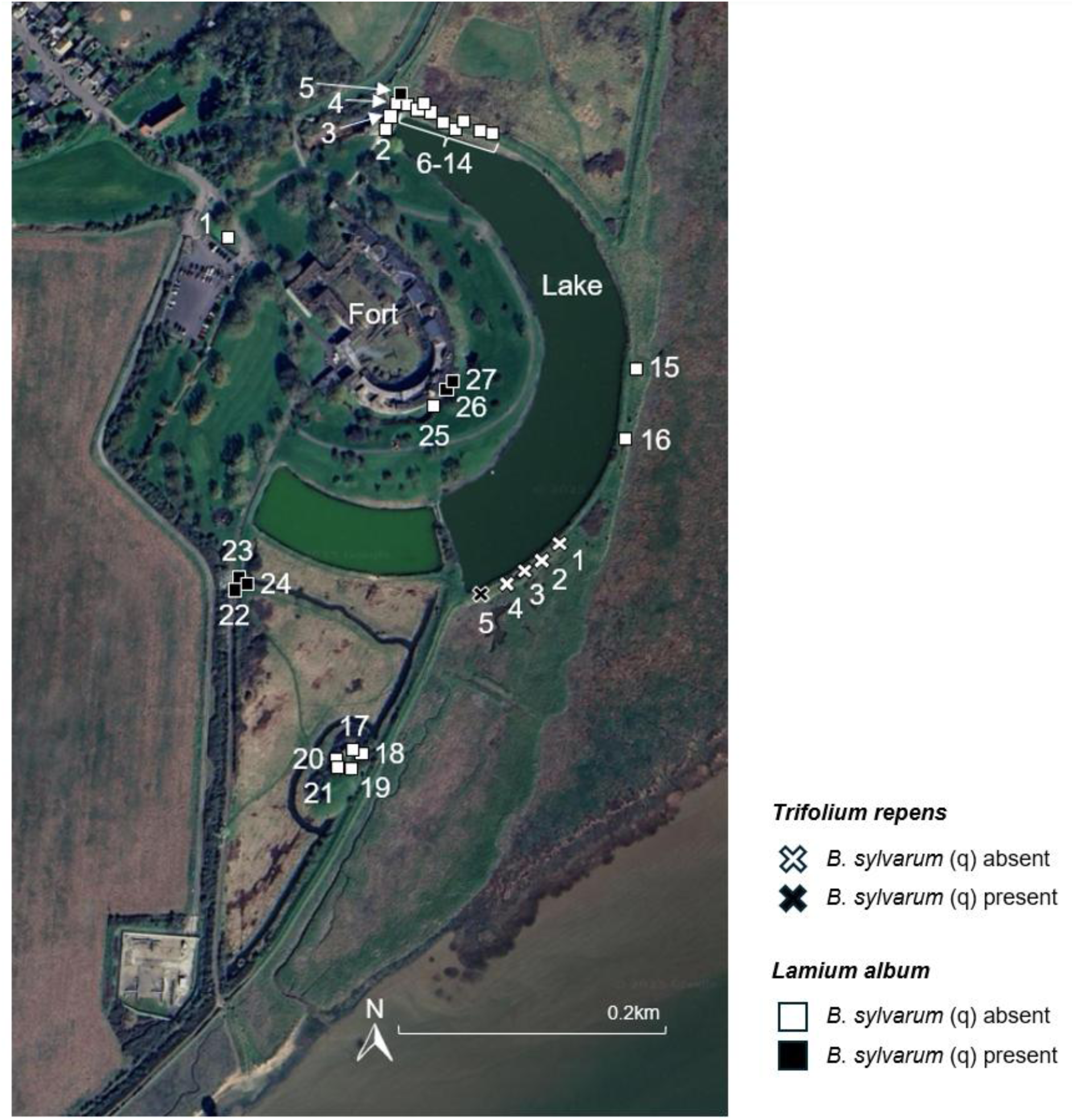
The distribution of 27 *Lamium album* (squares) and five *Trifolium repens* (crosses) patches around the fort. Black squares and crosses indicate which patches were visited by *B. sylvarum* queens (Google Earth 2025)

#### Quantifying *B. sylvarum* queens foraging trips within patches

On days with conditions favourable for PoMS FIT counts (see above), patches of *L. album* and *T. repens* were observed for 10 minutes. If a queen arrived at the patch within those 10 minutes, we followed her on her foraging trip within the patch, using a clicker to count the number of flowers she visited. We followed each queen until she left the patch, even if this exceeded the 10-minute period. If a queen did not arrive at a patch within 10 minutes, we moved onto another patch and repeated this procedure. These observations were carried out across 4 days, for a total of 97 observations. On the first day, the *L. album* and *T. repens* patches visited were those where *B. sylvarum* queens were seen foraging during the 5-minute observations made in the previous week. On the following three days of 10-minute observations, all accessible patches were observed (Fig. 2). Patches 9 and 14 had stopped flowering by the time we started the 10-minute observations of patches.

### Data collection in the SSSI grassland meadows

Data collection in the SSSI grassland took place between May and June 2025, once the meadows had started to flower, and *B. sylvarum* queens were no longer observed around the fort park area. Surveys were carried out between 09:00 and 17:30 GMT, when temperatures were above 15°C if it was cloudy, and above 13°C if it was sunny (see above).

#### Plant–pollinator network transects

Transect walks in the SSSI grassland meadows began at the end of May 2025, when plants in the meadows started flowering. All data were collected between 09:00 and 17:30 GMT. Although there are six meadows at this site, we only walked transects on four of these meadows. Meadow 3 was not flowering sufficiently for data collection, and Meadow 1 had nesting skylarks and flowering orchids that could easily be trampled, and, as it is a protected SSSI grassland, sampling was not permitted (Fig. 1c) (Thurrock Council 2013). The distribution of flowering plants determined whether transects were linear or non-linear. Transects were linear when the distribution was relatively homogeneous (Meadow 2). Transects were non-linear when the distribution was highly heterogeneous (Meadows 4, 5, 6). Both types of transect were 50 m long. We walked transects at 10 m per minute on each meadow and recorded any pollinator visits within 1m either side of the transect (Banaszak 1980; Connop 2007). We walked a total of 136 transects between the end of May 2025, and the end of June 2025. At least 32 transects were walked on each meadow, and at least once per day. Plants and insect flower visitors were identified as described above.

We estimated floral abundances for each of the four transects in 1 m² quadrats every five metres along each transect, resulting in ten abundance sub-samples per transect. These estimates were only repeated on days when there was a visible change in species composition and abundance. *Rhinanthus minor* and *Galega officinalis* floral abundances were estimated by counting each multi-flowered stem as one floral unit. *Vicia villosa* and *Lotus corniculatus* floral abundances were estimated by counting partial inflorescences (a group of flowers on one stem) as one floral unit. The remaining floral abundances were estimated by counting each flower head as a single floral unit. We acknowledge that estimating floral abundances without counting individual flowers for each species may over- or underestimate floral abundances, but counting individual flowers was not practically feasible.

### Data analyses

We conducted all data analyses using R version 4.3.2 (R Core Team 2023). Details for specific R packages used are provided below.

#### Floral preferences and foraging behaviour within patches

To determine whether queens had a floral preference across the nine species flowering in the park areas around the fort, a chi-square test was performed to see whether plant resource was significantly associated with *B. sylvarum* queen presence. To test whether queen foraging behaviour was influenced by patch size, we used Spearman’s rank correlations to test whether there were relationships between: the number of flowers visited per patch and the total number of flowers in a patch, and the percentage of flowers visited per patch and the total number of flowers in a patch.

#### Plant–pollinator networks

Using the transect data, we constructed a plant–pollinator bipartite network to explore potential competition for plant resources. The edges in the bipartite network corresponded to the interactions (flower visits) between the plants (first node) and pollinators (second node). The weight of each edge reflected the total number of interactions between each plant and pollinator. The plotweb() function was used to create a bipartite graph, and visweb() function to create an adjacency matrix. Plant abundance bars were added to the bipartite network graph manually using Microsoft PowerPoint (version 2508). Connectance and weighted nestedness were calculated using the networklevel() function. Connectance describes the possible interactions in the network and whether the network is robust to species loss, while weighted nestedness identifies whether specialists interact with a subset of the species interacted with by generalists, and the frequency of these interactions. Species specificity indexes (SSI) were calculated using the specieslevel() function. This measure identifies whether each pollinator species in the network is a specialist or generalist, in terms of plant resources foraged on. SSI values of 0 indicate a generalist species, interacting with many plant resources, while values of 1 indicate a specialist species, interacting exclusively with one resource. These metrics were calculated using the bipartite package in R (Dormann et al. 2009).

#### Network null models to test bumblebee plant preference and avoidance

The relative frequency of resource utilisation displayed by foraging animals is often used as a proxy for the relative importance of different resources at the species–population level. However, without accounting for the relative abundance of different resources within their environment, we cannot delineate passive resource selection (e.g., resource utilisation driven by the relative abundance and detectability of resources) and active resource selection (e.g., strong preference or avoidance of a resource by foragers) (Minnaar et al. 2015).

To determine whether different bumblebee species showed different patterns of relative resource preference or avoidance, we integrated both flower abundance data (per plant species) and bumblebee visit data from transects. Using these data, we computed null models to estimate relative resource utilisation for each pollinator species group under the null hypothesis that there were no resource preferences (i.e., if relative flower abundance was the only determinant of foraging with no active resource selection displayed by pollinators). We ran 1,000 null model simulations per bumblebee– plant-species pair to generate a predicted visit frequency estimate for each pair with 95% confidence intervals. When actual visit frequencies for a bumblebee–plant-species pair fell outside of the null-model-predicted 95% confidence interval, we interpreted this an indication of significant preference for (above upper limit), or avoidance (below lower limit) of, a specific plant species. In this context, preference includes switching to a resource with less interspecific resource competition. Similarly, avoidance includes pollinators avoiding a resource due to high interspecific resource competition. Since the only passive resource selection factor included in the null model is floral abundance, the model inherently assumes that floral units from different species are equally detectable by bumblebees (Minnaar et al. 2015) and that floral units from different species (as counted by us) are a biologically realistic representation of floral abundance from an individual bumblebee forager perspective. These assumptions are likely to be violated to some degree, so we interpret the null model results with caution.

Null models were computed and outputs plotted using the econullnetr package (Vaughan et al. 2018).

## Results

### *Bombus sylvarum* queens forage preferentially on *L. album* and *T. repens* after emergence from hibernation

Nine plant species flowered over the study season around the fort (Table 1). We conducted between 16 and 27 five-minute observations of pollinator visits to each flowering species (Table 1). We recorded twenty-three pollinator groups interacting with the flowers of these plants (Supplementary Table 1). Ten *B. sylvarum* queens, and three *B. sylvarum* workers were recorded across 1,050 minutes of observation. Queens were observed nine times on *L. album* flowers, and once on *T. repens. B. sylvarum* workers foraged on both *L. album* and *T. repens* flowers. There was a significant association between plant species and *B. sylvarum* queen presence (χ²_8_ = 55.18, *df* = 8, *p* = 4.076 x 10^-9^), which reflects the fact that *B. sylvarum* queens were almost only seen foraging on *L. album*.

**Table 1.** The number of *Bombus sylvarum* queen observations on the nine different plant species flowering around the fort in May 2025.

| Plant species | Total time plant species observed (minutes) | Number of *B. sylvarum* queens observed | Total number of observation periods |
| --- | --- | --- | --- |
| *Anthriscus sylvestris* | 135 | 0 | 27 |
| *Bellis perennis* | 130 | 0 | 26 |
| *Crataegus monogyna* | 135 | 0 | 27 |
| *Lamium album* | 135 | 9 | 27 |
| *Lepidium draba* | 135 | 0 | 27 |
| *Ranunculus acris* | 135 | 0 | 27 |
| *Taraxacum officinale* | 110 | 0 | 22 |
| *Tragopogon porrifolius* | 55 | 0 | 11 |
| *Trifolium repens* | 80 | 1 | 16 |

### Abundance of floral resources

There were 27 *L. album* patches (total floral units counted from 26 patches = 17,255) and five *T. repens* patches (total floral units = 330) around the fort (Fig. 2). *L. album* patch 4 was inaccessible, so its patch size and total floral units were not calculated (Fig. 3). The patch size of *L. album* patches 3 and 5 were not recorded due to human error (Fig. 3). The mean *L. album* patch size was 1.67 m² ± 1.91 (SD), and the mean *T. repens* patch size was 1.53 m² ± 1.31. The mean *L. album* floral units per patch was 664±1147, and the mean *T. repens* floral units per patch was 66±55.

**Fig. 3.**
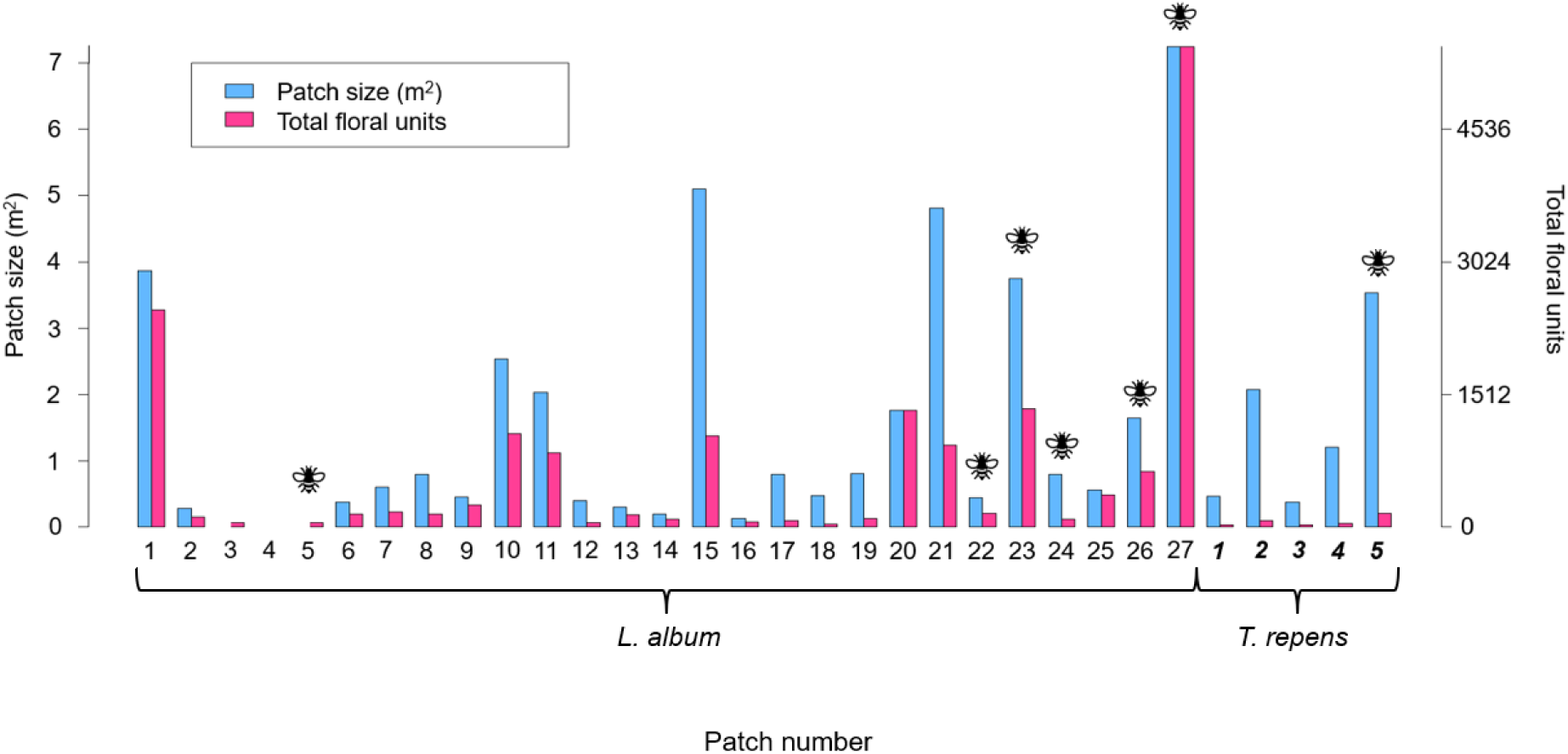
The patch sizes (m^2^) and total number of floral units for each of the *L. album* and *T. repens* (in bold and italic) patches around the fort. Patches visited by *B. sylvarum* queens are labelled with a bee icon. *L. album* patch 4 was inaccessible for calculating patch size and floral units. The size of *L. album* patches 3 and 5 were missed due to human error. Patch numbers refer to the numbered patches in Fig. 2 (see also Supplementary Table 2)

### Queen behaviour within patches

During the 10-minute observations, *B. sylvarum* queens were recorded most often at a temperature of 18°C (5/9 observations, with 1 at 16 °C, 2 at 17 °C, and 2 at 21 °C) and during occasional winds (5/9 observations, with 3 at gentle winds, and 2 at strong wind conditions). *B. sylvarum* queens were seen most often at patches on the lake side closer to the fort, where wind levels tended to be less strong than on the side of the lake further from the fort (Supplementary Table 3).

Since there was only one data point for *T. repens,* the following analyses were based on *L. album* patches. *B. sylvarum* queens visited approximately seven flowers per minute within patches (Supplementary Table 4). There was no relationship between the number of flowers visited and the total number of flowers in a patch (Spearman’s ρ = 0.590, *p* = 0.123) (Fig. 4a). In contrast, queens visited a significantly smaller percentage of flowers per patch as the total number of flowers per patch increased (Spearman’s ρ = -0.916, *p* = 0.001) (Fig. 4b). Patch area was significantly positively associated with total number of flowers per patch, suggesting that patch size is a good proxy for floral units (Spearman’s ρ = 0.926, *p* = 0.001) (Fig. 4c).

**Fig. 4.**
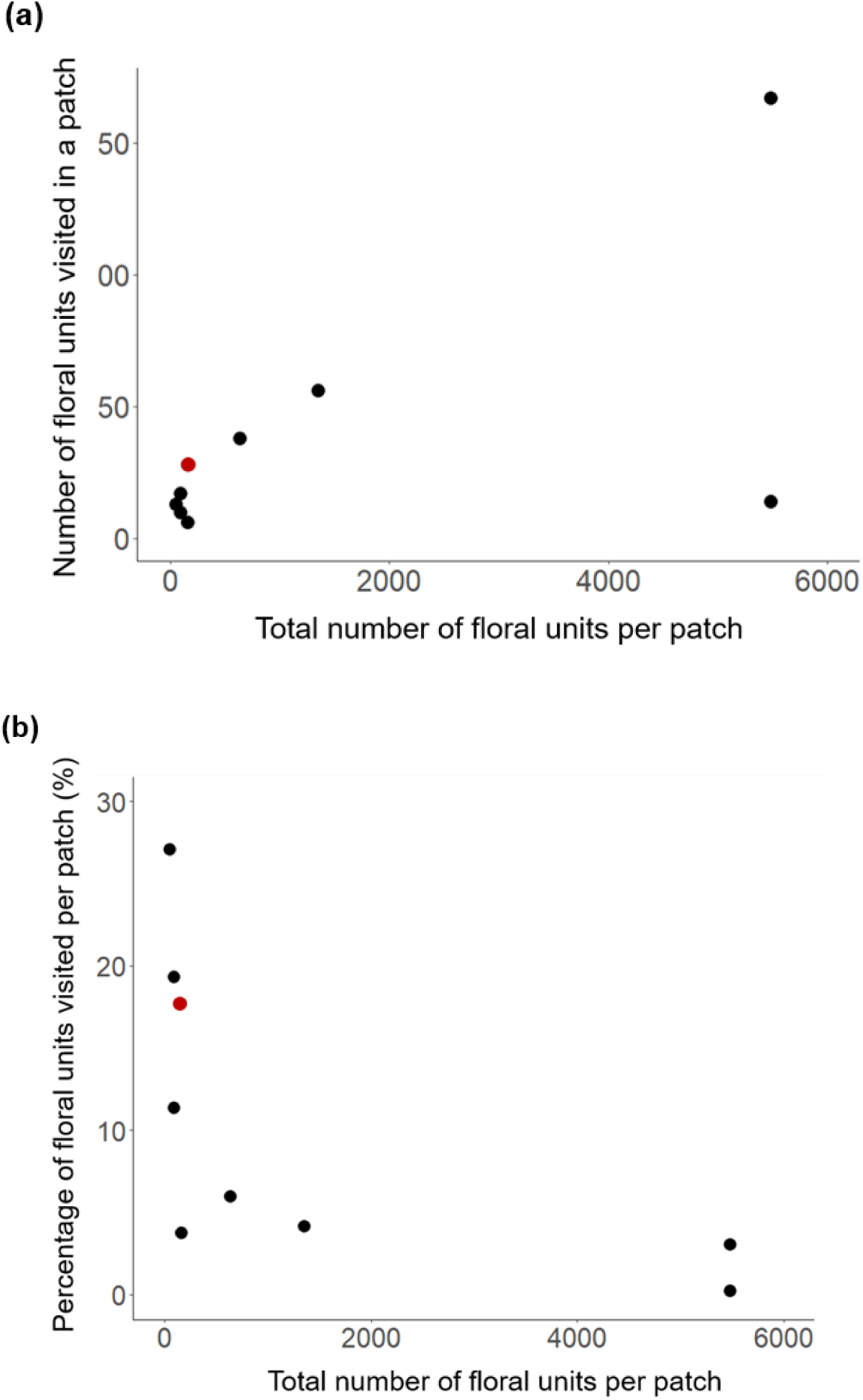
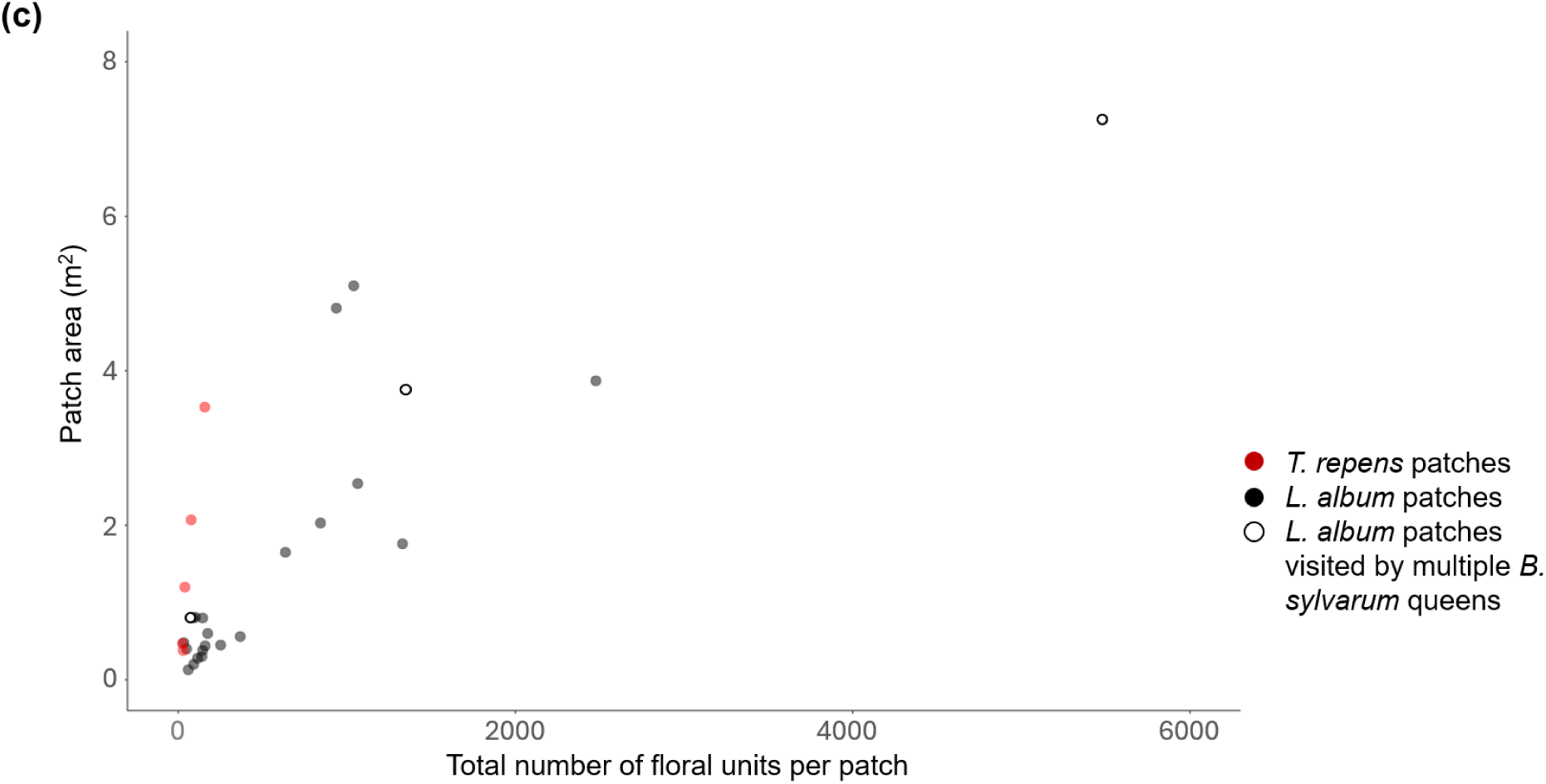
The **(a)** number of floral units visited; **(b)** percentage of floral units visited by *B. sylvarum* queens in *L. album* and *T. repens* patches around the fort, based on the number of floral units per patch; **(c)** area of each *L. album* and *T. repens* patch based on the number of floral units per patch. *L. album* patches are represented by black circles (*L. album* patches visited multiple times by *B. sylvarum* queens are shown as white circles). *T. repens* patches are represented by red circles. *L. album* patches 3-5 are not shown as they had some missing data (see Supplementary Table 2). While all analyses in the text are based on the *L. album* data, we have included the *T. repens* datapoints in the graphs for comparison

### Meadow plant–pollinator interactions

We recorded 2,049 plant–pollinator interactions between 27 flowering plant species, and 39 pollinator groups, across four meadows during May and June 2025 (Fig. 5; see also Supplementary Fig. 1 for separate plant–pollinator networks for each sampled meadow). *Lotus corniculatus, Rhinanthus minor, Scorzoneroides autumnalis, Trifolium pratense,* and *Vicia villosa* flowered across the duration of the whole season (Supplementary Table 5). *Scorzoneroides autumnalis* (363) and *Vicia villosa* (607) had the greatest number of interactions with pollinators (seen as the darkest cells in Fig. 6; see also Supplementary Fig. 2 for separate adjacency matrixes for each sampled meadow). *Bombus terrestris/ lucorum* workers (459) and *Coleoptera d* (358) had the greatest number of interactions with plants (Fig. 5,6). The connectance value was low for the overall network (0.181), which indicates that the number of plant–pollinator interactions observed was lower than the total number of possible interactions (Supplementary Table 6). The weighted nestedness was very low (0.456), which means that specialists and generalists did not have much overlap between the species they visited (Supplementary Table 6). Fifteen pollinator species had species specificity indexes of one (total specialists), with eight of these species only recorded once over the study season.

**Fig. 5.**
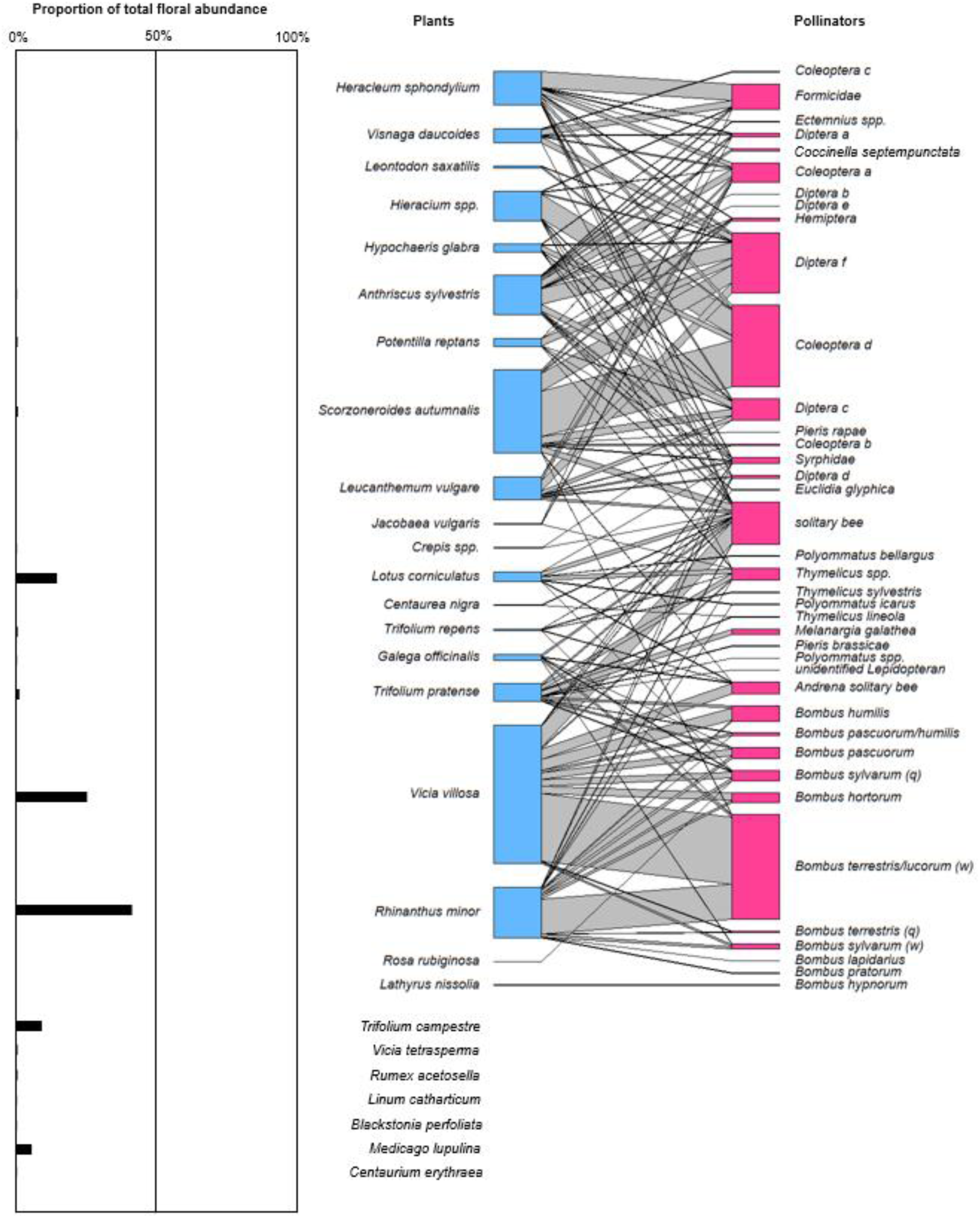
(right) The interaction network between plants and pollinators recorded during transect walks at Coalhouse Fort, Essex, in May-June 2025; ‘q’ = bumblebee queens; ‘w’ = bumblebee workers; (left) the abundance of each plant species in the network: some plant species do not have abundance measures as they were not found in the quadrats along each transect

**Fig. 6.**
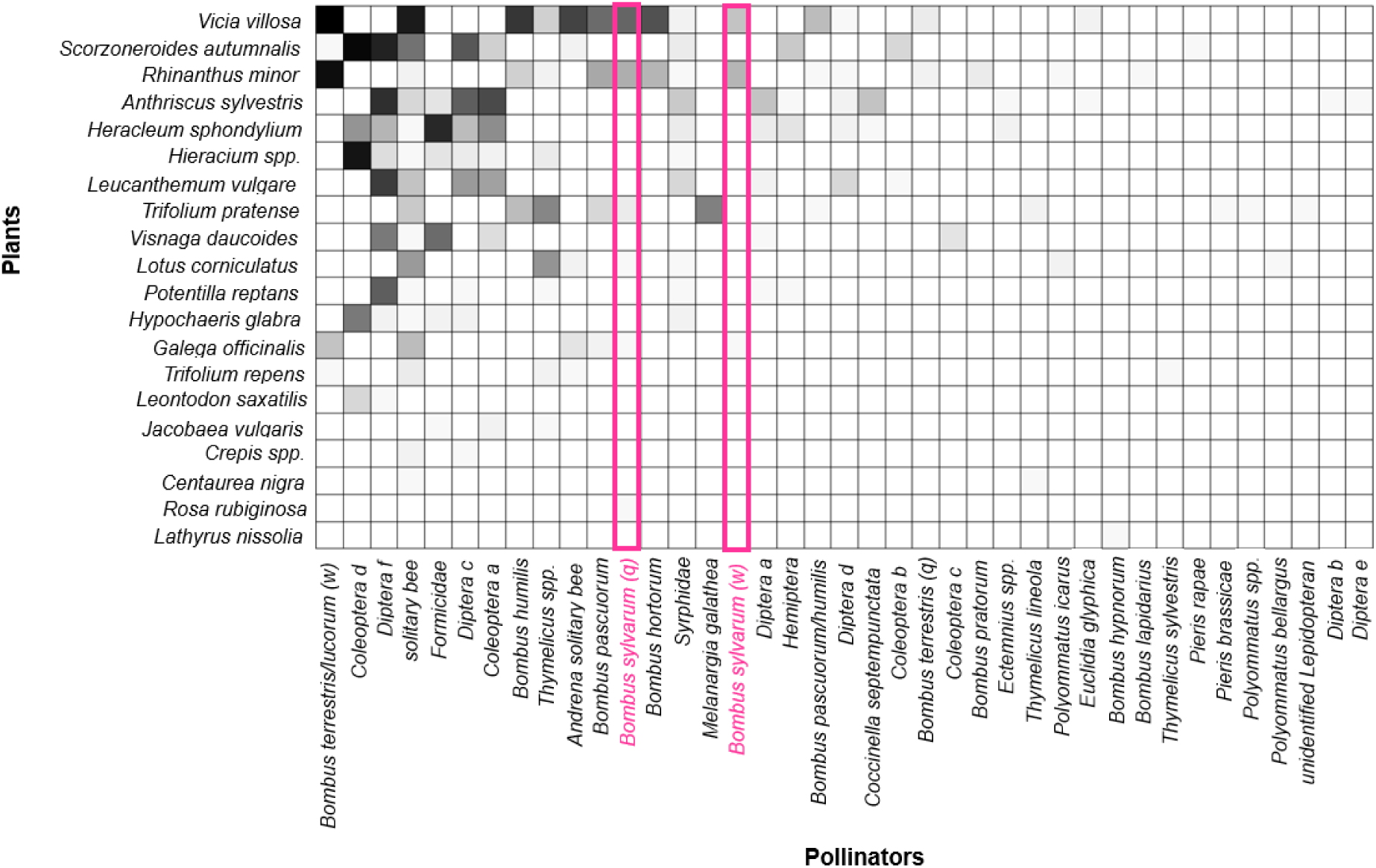
Adjacency matrix for the plants and pollinators seen interacting during the transect walks at Coalhouse Fort between May and June 2025. The darker the cell, the greater the number of interactions between the two species. Empty cells are indicative of no interactions. *Bombus sylvarum* (queen (‘q’) and worker (‘w’)) interactions are highlighted

*Bombus sylvarum* queens foraged on *V. villosa* (29)*, R. minor* (12)*, T. pratense* (3)*, Galega officinalis* (1)*, L. corniculatus* (1), and *Rosa rubiginosa* (1). In fact, the *B. sylvarum* queen foraging on *R. rubiginosa* was the only pollinator observed to have foraged on this plant species (Fig. 5). *B. sylvarum* workers also foraged on *R. minor* (11)*, V. villosa* (9), and *G. officinalis* (1). There is overlap between the plant species *B. sylvarum, B. terrestris/ lucorum* workers, and *Bombus humilis* foraged on (see bottom of Fig. 5 pollinator section). *B. terrestris/ lucorum* workers foraged on *V. villosa* (295)*, R. minor* (153)*, G. officinalis* (9)*, S. autumnalis* (1), and *T. repens* (1)*. B. humilis* foraged on *V. villosa*(50), *T. pratense* (10), and *R. minor* (7).

From the three bumblebee species mentioned above, *B. humilis* had the highest species specificity index (SSI = 0.754) in the full plant–pollinator network. *B. terrestris/ lucorum* workers had a slightly higher SSI value (0.707), than *B. sylvarum* queens (SSI = 0.650) and workers (SSI = 0.657).

#### Network null models to test bumblebee plant preference and avoidance

*Vicia villosa* and *T. pratense* were visited more often than expected by *B. sylvarum* queens, while *R. minor* and *L. corniculatus* were visited less often than expected by *B. sylvarum* queens, given the floral abundances available in the meadows (Fig. 7a). All bumblebees foraged on *R. minor* less than expected given its floral abundance in the meadows (Fig. 7). *B. sylvarum* workers visited *G. officinalis* more often than expected (however, with only one visit recorded, we do not consider this to be a robust result) (Fig. 7b). *B. humilis* visited *V. villosa* and *T. pratense* more often than expected (Fig. 7c). *Bombus pascuorum* visited *V. villosa, T. pratense,* and *G. officinalis* more often than expected (Fig. 7d). *Bombus terrestris/ lucorum* workers visited *V. villosa* and *G. officinalis* more often than expected (Fig. 7e).

**Fig. 7.**
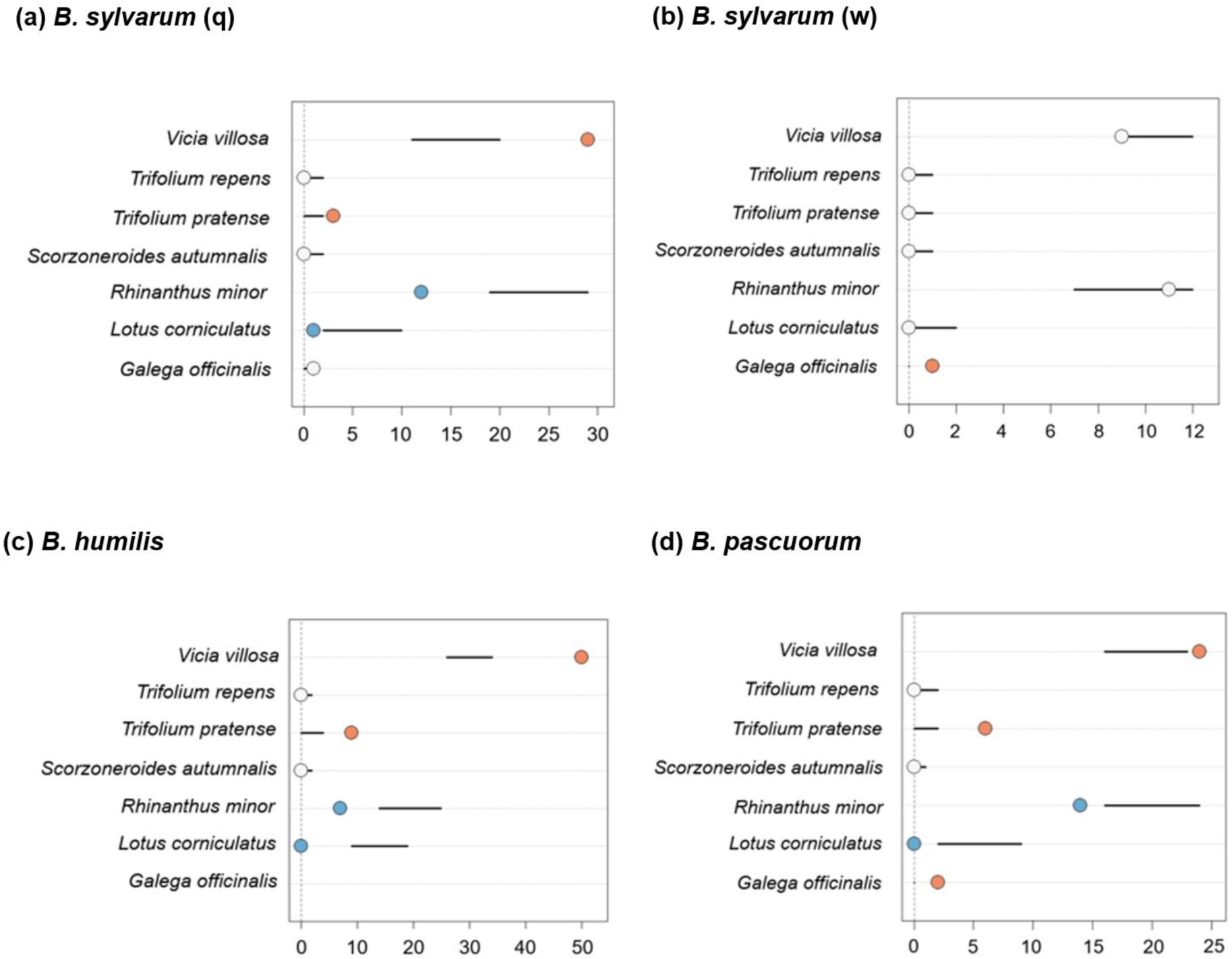
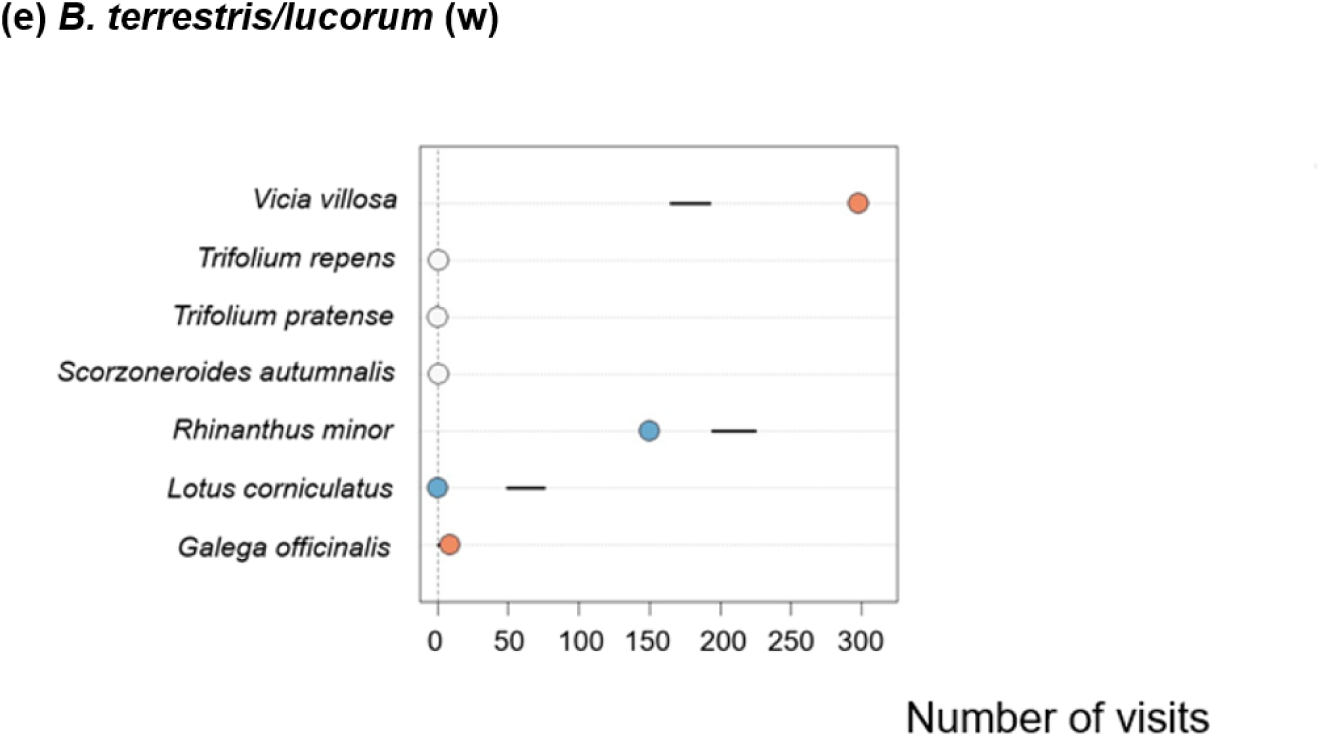
The number of visits to each plant species in comparison to null-model predicted confidence intervals for number of visits based on floral abundance for **(a)** *Bombus sylvarum* (q), **(b)** *Bombus sylvarum* (w), **(c)** *Bombus humilis,* **(d)** *Bombus pascuorum*, **(e)** *Bombus terrestris/ lucorum* (w). The lines on each plot represent the 95% confidence interval of the null model predictions of the number of visits. The actual number of visits to each plant is shown as a dot (blue dot: actual number of visits falls below the 95% confidence interval; white dot: actual number of visits falls within the 95% confidence interval; orange dot: actual number of visits is above the 95% confidence interval)

## Discussion

Early in the season, when a limited number of floral species were available, *B. sylvarum* queens foraged almost exclusively on *L. album.* The percentage of *L. album* flowers visited per patch significantly decreased with the total number of flowers per patch. Later in the season, when other plant species started flowering, *B. sylvarum* queens foraged on a broader array of species, with a preference for *T. pratense* and *V. villosa.* Queens shared many of these resources with *B. sylvarum* workers, *B. terrestris/ lucorum* workers, and *B. humilis,* and had a slightly lower specificity index than other bumblebee species. This suggests that *B. sylvarum* queens are not more specialised in their resource utilisation relative to workers from other bumblebee species and are likely facing significant competition from them.

The *B. sylvarum* queens at Coalhouse Fort were first seen on 15^th^ April 2025 (Reeves 2025, pers. comm.). The typical emergence time of *B. sylvarum* queens is late April or May, but this has become earlier in recent years (as seen in this study and in Horsley 2024), possibly due to climate change (Benton 2006; Horsley 2024). *B. sylvarum* queens were only seen foraging on *L. album* and *T. repens*around the fort. Both these plant species have flowers with deep or narrow corollas. While *B. sylvarum* queens were only seen foraging on *T. repens* once, all the *T. repens* patches were found on the outer edge of the lake. Interestingly, the *L. album* patches that were found on that side of the water were also not visited by *B. sylvarum* queens. It seems likely that the generally stronger winds on the outer edge of the lake resulted in the lack of foraging visits to the deep-flowered plant species found there, since *B. sylvarum* queens were most often seen during observation periods when leaves were moving occasionally or gently all the time around the fort (UK Pollinator Monitoring Scheme 2025).

We found a negative correlation between total number of floral units in a patch and the percentage of floral units visited per patch by *B. sylvarum* queens. This is expected since bumblebees tend to visit fewer floral units in patches as patch size increases (Goulson 2000). The maximum percentage of floral units visited in a single patch was 27.1%, suggesting that the number of *L. album* floral units may not have been a limiting resource. However, to determine whether there is a sufficient abundance of *L. album* at a given site for *B. sylvarum* queens at this time of year, we must understand how much nectar, and therefore sugar content, these flowers produce, and the daily nutritional requirements of queens.

Individual *L. album* flowers live for an average of 3–4 days, before the corolla starts wilting (Sulborska et al. 2014; Sulborska-Różycka et al. 2023). The average sugar mass per *L. album* flower is 1.08 mg, and the average sugar content in *L. album* nectar is 43.87% (calculated using data from: Jabłoński 1960; Gulyás 1967; Maurizio and Grafl 1969; Sulborska et al. 2014; Baude et al. 2016). A sucrose concentration of 50-65% in nectar maximises the net energy uptake for bees because they lap the nectar (dipping tongue in and out) (Harder 1986; Nardone et al. 2013). This suggests that *L. album* flowers are a relatively good nectar resource for *B. sylvarum.* The rate of foraging for *B. sylvarum* queens was calculated to be about seven flowers per minute. With an average patch size of 1.67 m^2^, and 664 flowers, the average number of *L. album* flowers in 1 m^2^ was 398. Using the average sugar mass per *L. album* flower, we can calculate the average sugar mass per m^2^ of *L. album* flowers: 430 mg. A honeybee worker requires 11 mg of dry sugar a day (Tsuruda et al. 2021). While the requirements for a bumblebee queen may be greater than this, the sugar mass from a m^2^ patch of *L. album* is enough to support 39 honeybee workers, and likely a good number of *B. sylvarum* queens. Although *L. album* and *T. repens* resources were shared with other bumblebees and pollinators, *B. sylvarum* queens were the main visitors. This suggests that early floral resources are currently unlikely to be limiting to *B. sylvarum* queens at Coalhouse Fort.

The management plan at Coalhouse Fort involves regular mowing of the grass around the publicly accessible fort park area (Thurrock Council 2013), using a ride-on mower and hand-held grass trimmer (pers. obs.). To maximise the foraging quality of this, and other sites for endangered *B. sylvarum* queens, mowing should be reduced around *L. album* patches to allow these patches to spread. *L. album* abundance can also be increased by seeding areas of bare soil (RHS 2025). In addition, as *B. sylvarum* queens were most often seen at lower wind levels, management to enhance the success of *B. sylvarum* spring queens should focus on maintaining and growing populations of *L. album* in wind-sheltered areas.

While *Bombus sylvarum* queens were initially restricted to the two plant species with deep corollas flowering around the fort (*L. album* and *T. repens*), when meadows started flowering, *B. sylvarum* queens had access to five floral species with deep or narrow corollas. However, *B. sylvarum* queens only foraged on four of these plant species. Surprisingly, *B. sylvarum* queens did not forage on *T. repens* in the meadows, but did forage on its congener, *T. pratense*, and more often than expected by null models*. T. pratense* has a higher nectar sugar content (116.86 μg per flower per day) than *T. repens* (48.97) (Baude et al. 2016), which could explain this preference. In addition to its high nectar sugar content, *T. pratense* may be favoured by long-proboscid pollinators as a resource because of its deep and narrow corolla tube (depth: 9.31 ± 0.41 mm; corolla opening width: 1.95 ± 0.07 (Sun et al. 2026)), which may be inaccessible to shorter-tongued competitors. Two other long-tongued bumblebees, *B. humilis* (8.1 ± 0.5 mm) and *B. pascuorum* (8.5 ± 0.6 mm) (Goulson et al. 2005), also foraged on *T. pratense* more often than expected by null models, and other visitors to *T. pratense* consisted mostly of long-proboscid butterflies.

In contrast, 14 and 15 pollinators foraged on *V. villosa* and *R. minor,* respectively, and these two species were dominated by visits from *B. terrestris/ lucorum* workers. While *V. villosa* has a deep corolla (11.64 ± 0.26 mm), its corolla is also relatively wide (2.83 ± 0.06 mm) (Sun et al. 2026). Easier access provided by a wider corolla may explain why short-tongued *B. terrestris/ lucorum* workers visited *V. villosa* more often than predicted by null models, while not visiting *T. pratense* at all. *T. pratense* may therefore be a valuable resource for long-tongued bumblebees like *B. sylvarum* and *B. humilis* (UK BAP species (UKBAP 2007)) because of reduced resource competition with highly abundant *B. terrestris/ lucorum* workers. Maintaining *T. pratense* in meadows may therefore be beneficial for improving *B. sylvarum* queen abundance.

However, despite more intense resource competition on *V. villosa*, *B. sylvarum* queens also visited this species more often than predicted by null models. The mean nectar sugar content of *V. villosa* is 166 μg per flower (Stpiczyńska and Pielecki 1999), making it a potentially valuable resource for *B. sylvarum* queens despite the potential for resource competition. Short-tongued *Bombus terrestris/ lucorum* workers appeared to be more specialised, or limited, in resource use than *B. sylvarum* queens and workers. *B. sylvarum* queens visited the highest number of plant species in the meadow network, compared to other bumblebees (6 of 27). A similar observation was made by Williams (2005) at Dungeness in Southern England. Since the *B. sylvarum* queen interactions (47) were much less common than the *B. terrestris/ lucorum* worker interactions (459), but *B. sylvarum* queens still visited the highest number of plant species (compared to other bumblebees), this difference in specialisation appears to be robust to sample size. Therefore, while maintaining key species like *T. pratense* may be beneficial for improving *B. sylvarum* queen abundance, managers should also ensure diversity in floral resources.

In conclusion, this study improves our understanding of foraging preferences and requirements for *B. sylvarum* queens. Since the queen stage of the bumblebee life cycle is essential for establishing new colonies, and increasing numbers, it is vital to protect and increase the abundance of plant species that *B. sylvarum* queens prefer to forage on. Our results particularly highlight the importance of *L. album* at the start of the foraging period for *B. sylvarum* queens, and site management to maximise populations of this species would clearly be advantageous. While our study is limited to a single site and, as is the nature of working with rare species, a limited number of observations, it provides the groundwork for quantifying the foraging preferences and needs of this and other rare bumblebee species at the start of the lifecycle. Conducting similar work at other sites known to support *B. sylvarum* queens will identify whether the foraging patterns at Coalhouse Fort are common or unique in relation to other populations, and this will inform decisions about management across sites to support populations of this iconic species.

## Acknowledgements

We would like to thank the Brown and Leadbeater lab groups for all their help. Thank you to Dr. Becky Thomas for all her support. Thank you also to Ray Reeves, the ranger at Coalhouse Fort, for sharing his knowledge and wisdom of the site and the shrill carder bumble bee. Thank you to Daisy Headley, at the Bumblebee Conservation Trust, for her help with coordinating the project. Many thanks to Professor Lynn Dicks and Jacqui James for their ongoing collaboration and support with the research.

## Author contributions

Mark JF Brown and Julia de Korab Skulska contributed to the study conception and design. Fieldwork, analysis, and the first draft of the manuscript was completed by Julia de Korab Skulska. Corneile Minnaar conducted the null model data analysis and wrote the method section for this. Mark JF Brown and Corneile Minnaar commented on previous versions of the manuscript. All authors read and approved the final draft of the manuscript.

## Corresponding author

Correspondence to Julia de Korab Skulska

## Ethics declaration

### Ethical approval

Ethical approval was obtained from Royal Holloway, University of London. No animals were harmed during this study.

### Competing interests

The authors have no competing interests.

### Data availability

Data and code will be made publicly available on an open-access repository (DRYAD) at the time of peer-reviewed publication.

## Notes

### Competing Interest Statement

The authors have declared no competing interest.

