## Supplementary Table 1 for "Quantitative evidence for foraging preferences of shrill carder bumble bee (*Bombus sylvarum*) queens"

**Supplementary Table 1** The frequency of each of the 22 pollinator groups (first column) interacting with the flowers of nine plant species (first row) around the fort for each of the five minute observation periods

| Pollinator group ↓ | *Ranunculus acris* | *Taraxacum officinale* | *Bellis perennis* | *Lamium album* | *Crataegus monogyna* | *Tragopogon porrifolius* | *Lepidium draba* | *Trifolium repens* | *Anthriscus sylvestris* | Total frequency |
|---|---|---|---|---|---|---|---|---|---|---|
| *Bombus sylvarum* (q) | 0 | 0 | 0 | 0.33 | 0 | 0 | 0 | 0 | 0 | 0.33 |
| *Bombus sylvarum* (w) | 0 | 0 | 0 | 0.07 | 0 | 0 | 0 | 0.13 | 0 | 0.20 |
| Small black fly | 0.22 | 0.09 | 0.23 | 0 | 0.22 | 0 | 0.67 | 0 | 0.15 | 1.58 |
| Big black fly | 0.04 | 0 | 0 | 0 | 0 | 0 | 0 | 0 | 0.04 | 0.08 |
| Medium black fly | 0 | 0 | 0 | 0 | 0.04 | 0 | 0.07 | 0 | 0 | 0.11 |
| *Andrena* solitary bee | 0.15 | 0.09 | 0.15 | 0.07 | 0.30 | 0 | 0.11 | 0 | 0 | 0.87 |
| Pollen beetle | 0.07 | 0 | 0 | 0 | 0 | 0 | 0 | 0 | 0 | 0.07 |
| *Formicidae* | 0.04 | 0.27 | 0 | 0 | 0 | 0 | 0 | 0 | 0.19 | 0.50 |
| *Bombus terrestris/ lucorum* (w) | 0 | 0 | 0 | 0.07 | 0.19 | 0 | 0 | 0 | 0 | 0.26 |
| *Bibionidae* | 0 | 0 | 0 | 0 | 0.11 | 0 | 1.11 | 0 | 4.33 | 5.55 |
| *Bombus pratorum* | 0 | 0 | 0 | 0.11 | 0 | 0.09 | 0 | 0 | 0 | 0.20 |

| | | | | | | | | | |
|---|---|---|---|---|---|---|---|---|---|
| *Bombus humilis* | 0 | 0 | 0 | 0.15 | 0 | 0 | 0 | 0 | 0 | 0.15 |
| *Syrphidae* | 0.11 | 0 | 0.04 | 0.04 | 0.15 | 0 | 0.04 | 0.06 | 0 | 0.44 |
| *Coccinella septempunctata* | 0 | 0 | 0 | 0 | 0 | 0 | 0.07 | 0 | 0 | 0.07 |
| *Bombus hortorum* | 0 | 0 | 0 | 0.19 | 0.07 | 0 | 0 | 0 | 0 | 0.26 |
| *Tritomegas sexmaculatus* | 0 | 0 | 0 | 0 | 0 | 0 | 0.04 | 0 | 0 | 0.04 |
| *Pieris rapae* | 0.04 | 0 | 0 | 0 | 0 | 0 | 0 | 0 | 0 | 0.04 |
| *Thymelicus* spp. | 0.04 | 0 | 0 | 0 | 0 | 0 | 0 | 0 | 0 | 0.04 |
| Shiny green beetle | 0 | 0 | 0 | 0 | 0 | 0 | 0.11 | 0 | 0 | 0.11 |
| *Bombus pascuorum* | 0 | 0 | 0 | 0.15 | 0 | 0 | 0 | 0 | 0 | 0.15 |
| Black beetle | 0.04 | 0 | 0 | 0 | 0 | 0 | 0.04 | 0 | 0 | 0.08 |
| *Glyphipterix simpliciella* | 0.04 | 0 | 0 | 0 | 0 | 0 | 0 | 0 | 0 | 0.04 |
