## Supplementary Table 2 for "Quantitative evidence for foraging preferences of shrill carder bumble bee (*Bombus sylvarum*) queens"

**Supplementary Table 2** The patch sizes ($m^2$) and total number of floral units for each of the *L. album* (above the thick line) and *T. repens* (below the thick line) patches around the fort. Patches visited by *B. sylvarum* queens are shown in bold. Patch numbers refer to the numbered patches in Fig. 3

| Patch number | Patch size (m²) | Total floral units |
|---|---|---|
| 1 | 3.87 | 2477 |
| 2 | 0.28 | 115 |
| 3 | NA | 48 |
| 4 | NA | NA |
| **5** | **NA** | **48** |
| 6 | 0.38 | 145 |
| 7 | 0.6 | 175 |
| 8 | 0.8 | 145 |
| 9 | 0.45 | 252 |
| 10 | 2.54 | 1065 |
| 11 | 2.03 | 844 |
| 12 | 0.4 | 50 |
| 13 | 0.3 | 140 |
| 14 | 0.2 | 92 |
| 15 | 5.1 | 1041 |
| 16 | 0.13 | 60 |
| 17 | 0.8 | 72 |
| 18 | 0.48 | 33 |
| 19 | 0.81 | 102 |
| 20 | 1.76 | 1330 |
| 21 | 4.81 | 938 |
| **22** | **0.44** | **160** |
| **23** | **3.75** | **1350** |
| **24** | **0.8** | **88** |
| 25 | 0.56 | 368 |
| **26** | **1.65** | **636** |
| **27** | **7.25** | **5481** |
| 1 | 0.47 | 25 |
| 2 | 2.07 | 77 |
| 3 | 0.38 | 30 |

| | | |
|---|---|---|
| 4 | 1.2 | 40 |
| **5** | **3.53** | **158** |
