## Supplementary Table 3 for "Quantitative evidence for foraging preferences of shrill carder bumble bee (*Bombus sylvarum*) queens"

**Supplementary Table 3** The number of *B. sylvarum* queens present at each patch of *L. album* (above the thick line) and *T. repens* (below the thick line) during 10-minute observations at different **(a)** temperatures; **(b)** wind levels. **(c)** the average wind level and temperature at each *L. album* and *T. repens* patch

**(a)**

| Patch number ↓ | 15°C | 16°C | 17°C | 18°C | 19°C | 20°C | 21°C |
|---|---|---|---|---|---|---|---|
| 1 | 0 | 0 | 0 | 0 | 0 | 0 | 0 |
| 2 | 0 | 0 | 0 | 0 | 0 | 0 | 0 |
| 3 | 0 | 0 | 0 | 0 | 0 | 0 | 0 |
| 4 | 0 | 0 | 0 | 0 | 0 | 0 | 0 |
| 5 | 0 | **1** | 0 | 0 | 0 | 0 | 0 |
| 6 | 0 | 0 | 0 | 0 | 0 | 0 | 0 |
| 7 | 0 | 0 | 0 | 0 | 0 | 0 | 0 |
| 8 | 0 | 0 | 0 | 0 | 0 | 0 | 0 |
| 9 | 0 | 0 | 0 | 0 | 0 | 0 | 0 |
| 10 | 0 | 0 | 0 | 0 | 0 | 0 | 0 |
| 11 | 0 | 0 | 0 | 0 | 0 | 0 | 0 |
| 12 | 0 | 0 | 0 | 0 | 0 | 0 | 0 |
| 13 | 0 | 0 | 0 | 0 | 0 | 0 | 0 |
| 14 | 0 | 0 | 0 | 0 | 0 | 0 | 0 |
| 15 | 0 | 0 | 0 | 0 | 0 | 0 | 0 |
| 16 | 0 | 0 | 0 | 0 | 0 | 0 | 0 |
| 17 | 0 | 0 | 0 | 0 | 0 | 0 | 0 |
| 18 | 0 | 0 | 0 | 0 | 0 | 0 | 0 |
| 19 | 0 | 0 | 0 | 0 | 0 | 0 | 0 |
| 20 | 0 | 0 | 0 | 0 | 0 | 0 | 0 |
| 21 | 0 | 0 | 0 | 0 | 0 | 0 | 0 |
| 22 | 0 | 0 | **1** | 0 | 0 | 0 | 0 |
| 23 | 0 | 0 | 0 | **1** | 0 | 0 | **1** |
| 24 | 0 | 0 | **1** | **1** | 0 | 0 | 0 |
| 25 | 0 | 0 | 0 | 0 | 0 | 0 | 0 |
| 26 | 0 | 0 | 0 | **1** | 0 | 0 | 0 |

| | | | | | | | |
|---|---|---|---|---|---|---|---|
| 27 | 0 | 0 | 0 | **2** | 0 | 0 | 0 |
| 1 | 0 | 0 | 0 | 0 | 0 | 0 | 0 |
| 2 | 0 | 0 | 0 | 0 | 0 | 0 | 0 |
| 3 | 0 | 0 | 0 | 0 | 0 | 0 | 0 |
| 4 | 0 | 0 | 0 | 0 | 0 | 0 | 0 |
| 5 | 0 | 0 | 0 | 0 | 0 | 0 | **1** |

**(b)**

| Patch number ↓ | Occasional | Gentle | Strong |
|---|---|---|---|
| 1 | 0 | 0 | 0 |
| 2 | 0 | 0 | 0 |
| 3 | 0 | 0 | 0 |
| 4 | 0 | 0 | 0 |
| 5 | **1** | 0 | 0 |
| 6 | 0 | 0 | 0 |
| 7 | 0 | 0 | 0 |
| 8 | 0 | 0 | 0 |
| 9 | 0 | 0 | 0 |
| 10 | 0 | 0 | 0 |
| 11 | 0 | 0 | 0 |
| 12 | 0 | 0 | 0 |
| 13 | 0 | 0 | 0 |
| 14 | 0 | 0 | 0 |
| 15 | 0 | 0 | 0 |
| 16 | 0 | 0 | 0 |
| 17 | 0 | 0 | 0 |
| 18 | 0 | 0 | 0 |
| 19 | 0 | 0 | 0 |
| 20 | 0 | 0 | 0 |
| 21 | 0 | 0 | 0 |
| 22 | 0 | 0 | **1** |

| | | | |
|---|---|---|---|
| 23 | **1** | **1** | 0 |
| 24 | **2** | 0 | 0 |
| 25 | 0 | 0 | 0 |
| 26 | 0 | **1** | 0 |
| 27 | **1** | **1** | 0 |
| 1 | 0 | 0 | 0 |
| 2 | 0 | 0 | 0 |
| 3 | 0 | 0 | 0 |
| 4 | 0 | 0 | 0 |
| 5 | 0 | 0 | **1** |

**(c)**

| Patch number | Average wind level | Average temperature (ºC) |
|---|---|---|
| 1 | strong | 18 |
| 2 | gentle | 17 |
| 3 | gentle | 17 |
| 4 | occasional | 16 |
| **5** | **occasional** | **16** |
| 6 | occasional | 18 |
| 7 | gentle | 17 |
| 8 | occasional/ gentle | 17 |
| 9 | occasional | 16 |
| 10 | occasional | 18 |
| 11 | occasional | 17 |
| 12 | gentle/ strong | 18 |
| 13 | occasional/ gentle | 18 |
| 14 | occasional | 17 |
| 15 | gentle/ strong | 18 |
| 16 | strong | 18 |
| 17 | gentle | 19 |
| 18 | strong | 18 |
| 19 | strong | 19 |
| 20 | gentle | 19 |
| 21 | gentle/ strong | 18 |

| | | |
|---|---|---|
| **22** | **occasional** | **19** |
| **23** | **occasional** | **19** |
| **24** | **occasional** | **19** |
| 25 | occasional/ gentle | 19 |
| **26** | **occasional** | **19** |
| **27** | **occasional** | **19** |
| 1 | strong | 18 |
| 2 | strong | 18 |
| 3 | strong | 18 |
| 4 | strong | 19 |
| **5** | **strong** | **19** |
