## Supplementary Table 4 for "Quantitative evidence for foraging preferences of shrill carder bumble bee (*Bombus sylvarum*) queens"

**Supplementary Table 4** The rate of *B. sylvarum* queen visitations in patches of *L. album* (above the thick line) and *T. repens* (below the thick line) as the number of flowers visited per minute. NA values are shown when the start or end time of a queen foraging trip was missed

| Patch number | Number of flowers visited | Foraging start time | Foraging end time | Foraging trip length (min) | Rate of foraging (flowers/min) |
|---|---|---|---|---|---|
| 5 | 13 | 09:58 | 10:02 | 4 | 3.25 |
| 22 | 6 | 16:17 | 16:19 | 2 | 3 |
| 23 | NA | 12:52 | 12:57 | 5 | NA |
| 23 | 56 | 16:06 | 16:13 | 7 | 8 |
| 24 | 17 | 13:10 | 13:11 | 1 | 17 |
| 24 | 10 | 11:01 | 11:03 | 2 | 5 |
| 26 | 38 | 15:22 | NA | NA | NA |
| 27 | 167 | NA | 15:35 | NA | NA |
| 27 | 14 | 11:29 | 11:31 | 2 | 7 |
| 5 | 28 | 14:39 | 14:44 | 5 | 5.6 |
