## Supplementary Table 5 for "Quantitative evidence for foraging preferences of shrill carder bumble bee (*Bombus sylvarum*) queens"

**Supplementary Table 5** Flowering plant species in the meadows at Coalhouse Fort, and the time periods when they were found during the 2025 study season

| Plant species | Area found | Time period |
|---|---|---|
| *Anthriscus sylvestris* | Meadow 4<br>Meadow 6 | 30$^{th}$ May – 24$^{th}$ June<br>30$^{th}$ May – 24$^{th}$ June |
| *Blackstonia perfoliata* | Meadow 2 | 8$^{th}$ – 24$^{th}$ June |
| *Centaurea nigra* | Meadow 6 | 25$^{th}$ – 27$^{th}$ June |
| *Centaurium erythraea* | Meadow 2<br>Meadow 5 | 18$^{th}$ – 27$^{th}$ June<br>25$^{th}$ – 27$^{th}$ June |
| *Crepis spp.* | Meadow 4<br>Meadow 5<br>Meadow 6 | 25$^{th}$ – 27$^{th}$ June<br>18$^{th}$ – 24$^{th}$ June<br>25$^{th}$ – 27$^{th}$ June |
| *Galega officinalis* | Meadow 6 | 13$^{th}$ – 27$^{th}$ June |
| *Heracleum sphondylium* | Meadow 6 | 8$^{th}$ – 24$^{th}$ June |
| *Hieracium spp.* | Meadow 5<br>Meadow 6 | 18$^{th}$ – 24$^{th}$ June<br>18$^{th}$ – 27$^{th}$ June |
| *Hypochaeris glabra* | Meadow 2<br>Meadow 4<br>Meadow 5 | 18$^{th}$ – 24$^{th}$ June<br>18$^{th}$ – 24$^{th}$ June<br>18$^{th}$ – 24$^{th}$ June |
| *Jacobaea vulgaris* | Meadow 6 | 18$^{th}$ – 24$^{th}$ June |
| *Lathyrus nissolia* | Meadow 4 | 30$^{th}$ May – 7$^{th}$ June |
| *Leontodon saxatilis* | Meadow 2 | 25$^{th}$ – 27$^{th}$ June |
| *Leucanthemum vulgare* | Meadow 6 | 30$^{th}$ May – 12$^{th}$ June |
| *Linum catharticum* | Meadow 2 | 30$^{th}$ May – 7$^{th}$ June |
| *Lotus corniculatus* | Meadow 4<br>Meadow 5<br>Meadow 6 | 30$^{th}$ May – 27$^{th}$ June<br>30$^{th}$ May – 27$^{th}$ June<br>13$^{th}$ – 27$^{th}$ June |
| *Medicago lupulina* | Meadow 2 | 13$^{th}$ – 27$^{th}$ June |
| *Potentilla reptans* | Meadow 6 | M, J1, J2, J3 |
| *Rhinanthus minor* | Meadow 2<br>Meadow 4<br>Meadow 5<br>Meadow 6 | 30$^{th}$ May – 27$^{th}$ June<br>30$^{th}$ May – 27$^{th}$ June<br>30$^{th}$ May – 27$^{th}$ June<br>30$^{th}$ May – 17$^{th}$ June |
| *Rosa rubiginosa* | Meadow 5 | 30$^{th}$ May – 7$^{th}$ June |
| *Rumex acetosella* | Meadow 2 | 30$^{th}$ May – 12$^{th}$ June |

| | | |
|---|---|---|
| *Scorzoneroides autumnalis* | Meadow 2<br>Meadow 4<br>Meadow 5 | 30th May – 27th June<br>30th May – 27th June<br>8th – 24th June |
| *Trifolium campestre* | Meadow 2<br>Meadow 4<br>Meadow 5<br>Meadow 6 | 30th May – 24th June<br>30th May – 24th June<br>8th – 24th June<br>30th May – 24th June |
| *Trifolium pratense* | Meadow 2<br>Meadow 4 | 8th – 27th June<br>30th May – 27th June |
| *Trifolium repens* | Meadow 4 | 30th May – 24th June |
| *Vicia tetrasperma* | Meadow 4<br>Meadow 6 | 30th May – 24th June<br>30th May – 7th June |
| *Vicia villosa* | Meadow 4<br>Meadow 5<br>Meadow 6 | 30th May – 24th June<br>30th May – 24th June<br>30th May – 27th June |
| *Visnaga daucoides* | Meadow 5<br>Meadow 6 | 25th – 27th June<br>25th – 27th June |
