## Supplementary Table 6 for "Quantitative evidence for foraging preferences of shrill carder bumble bee (*Bombus sylvarum*) queens"

**Supplementary Table 6** The connectance and weighted nestedness values for each of the four meadows sampled in the SSSI grassland

| Meadow number | Connectance | Weighted nestedness |
|---|---|---|
| 2 | 0.255 | 0.382 |
| 4 | 0.225 | 0.297 |
| 5 | 0.248 | 0.525 |
| 6 | 0.231 | 0.484 |
