## Supplementary Figure 1 for "Quantitative evidence for foraging preferences of shrill carder bumble bee (*Bombus sylvarum*) queens"

(a)

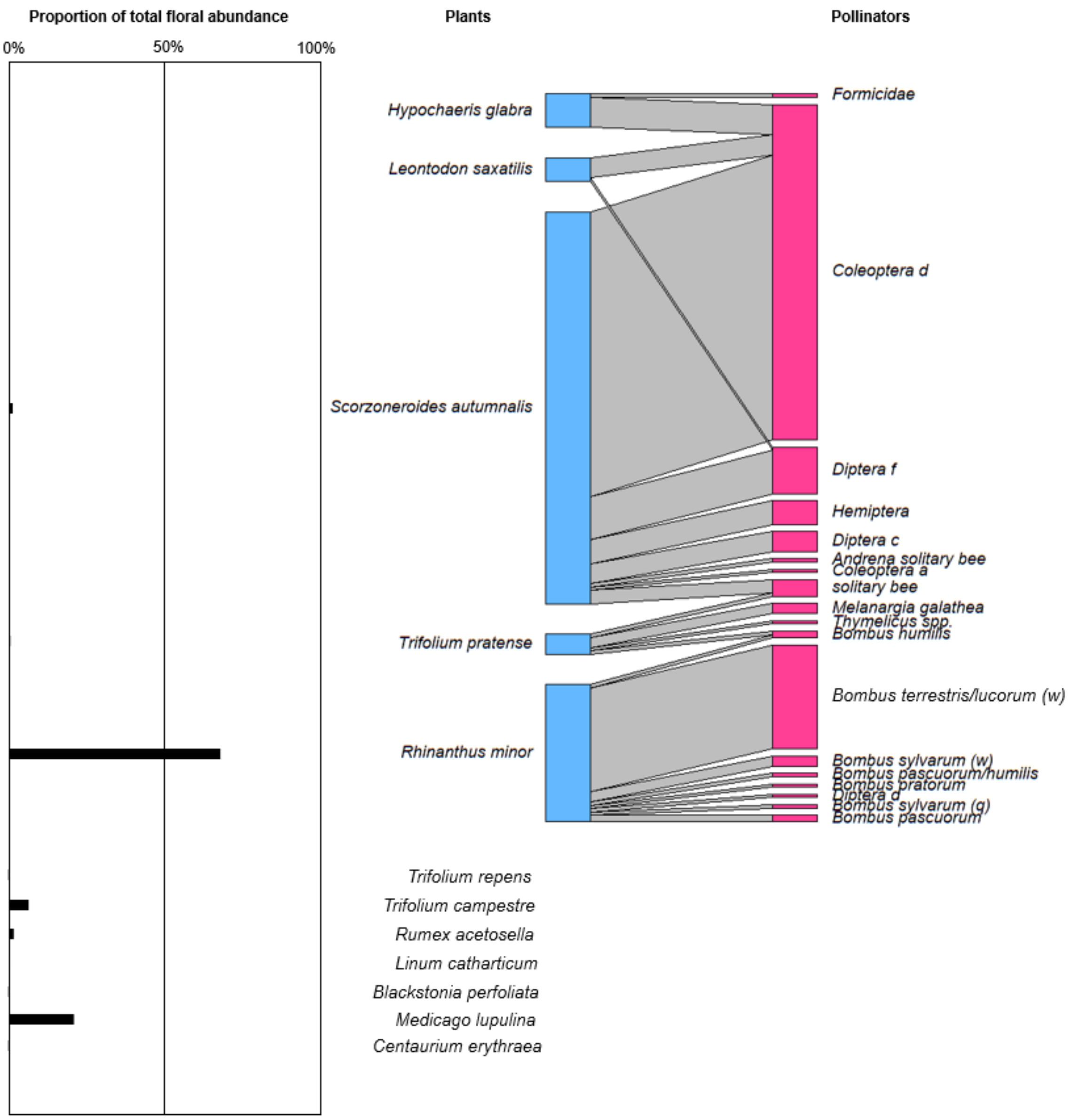
Proportion of total floral abundance
0%
50%
100%
Plants
Pollinators
Hypochaeris glabra
Leontodon saxatilis
Scorzoneroides autumnalis
Trifolium pratense
Rhinanthus minor
Trifolium repens
Trifolium campestre
Rumex acetosella
Linum catharticum
Blackstonia perfoliata
Medicago lupulina
Centaurium erythraea
Formicidae
Coleoptera d
Diptera f
Hemiptera
Diptera c
Andrena solitary bee
Coleoptera a
solitary bee
Melanargia galathea
Thymelicus spp.
Bombus humilis
Bombus terrestris/lucorum (w)
Bombus sylvarum (w)
Bombus pascuorum/humilis
Bombus pratorum
Diptera d
Bombus sylvarum (q)
Bombus pascuorum

(b)

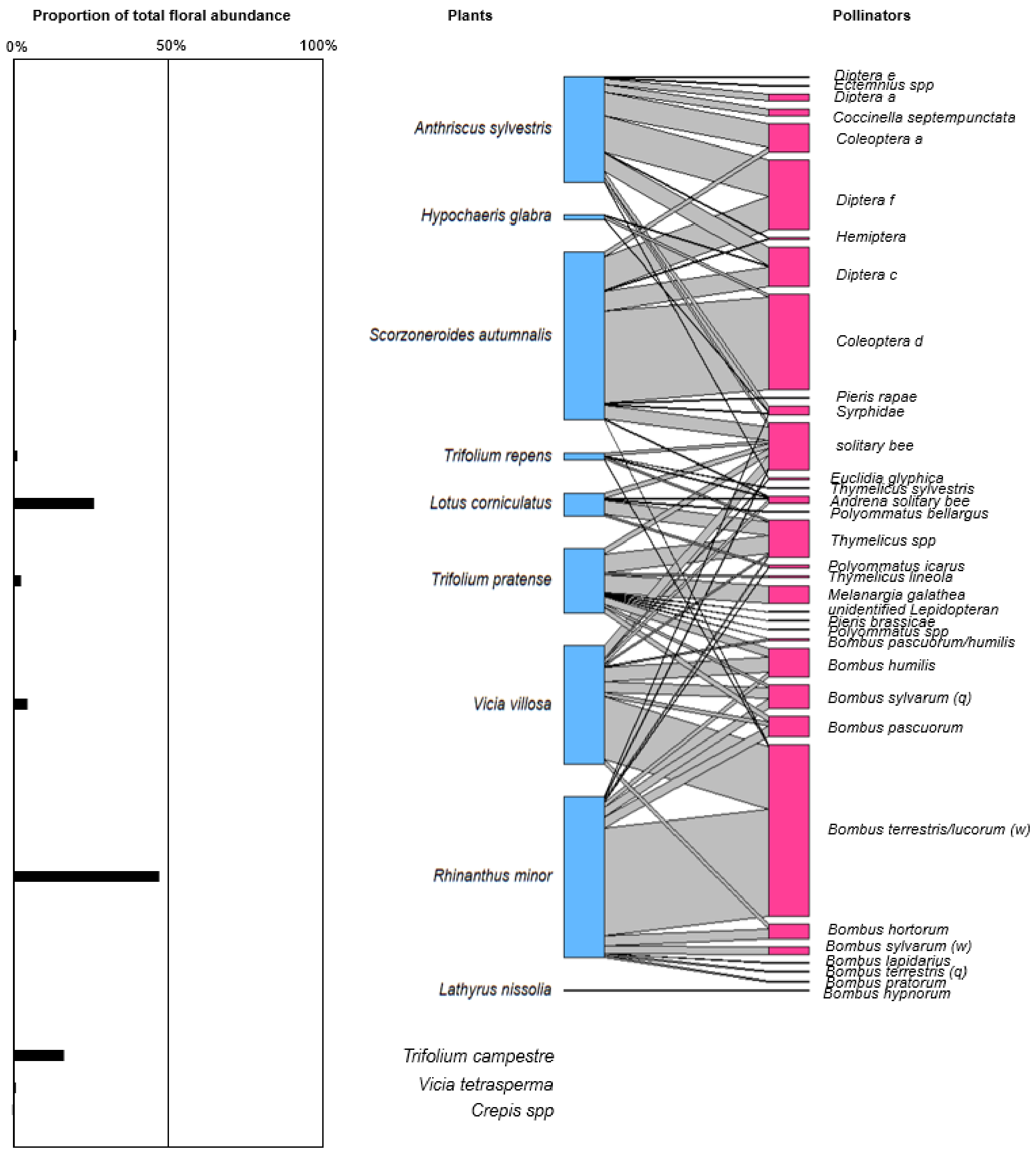
Proportion of total floral abundance
0%
50%
100%
Plants
Pollinators
Anthriscus sylvestris
Hypochaeris glabra
Scorzoneroides autumnalis
Trifolium repens
Lotus corniculatus
Trifolium pratense
Vicia villosa
Rhinanthus minor
Lathyrus nissolia
Trifolium campestre
Vicia tetrasperma
Crepis spp
Diptera e
Ectemnius spp
Diptera a
Coccinella septempunctata
Coleoptera a
Diptera f
Hemiptera
Diptera c
Coleoptera d
Pieris rapae
Syrphidae
solitary bee
Euclidia glyphica
Thymelicus sylvestris
Andrena solitary bee
Polyommatus bellargus
Thymelicus spp
Polyommatus icarus
Thymelicus lineola
Melanargia galathea
unidentified Lepidopteran
Pieris brassicae
Polyommatus spp
Bombus pascuorum/humilis
Bombus humilis
Bombus sylvarum (q)
Bombus pascuorum
Bombus terrestris/lucorum (w)
Bombus hortorum
Bombus sylvarum (w)
Bombus lapidarius
Bombus terrestris (q)
Bombus pratorum
Bombus hypnorum

(c)

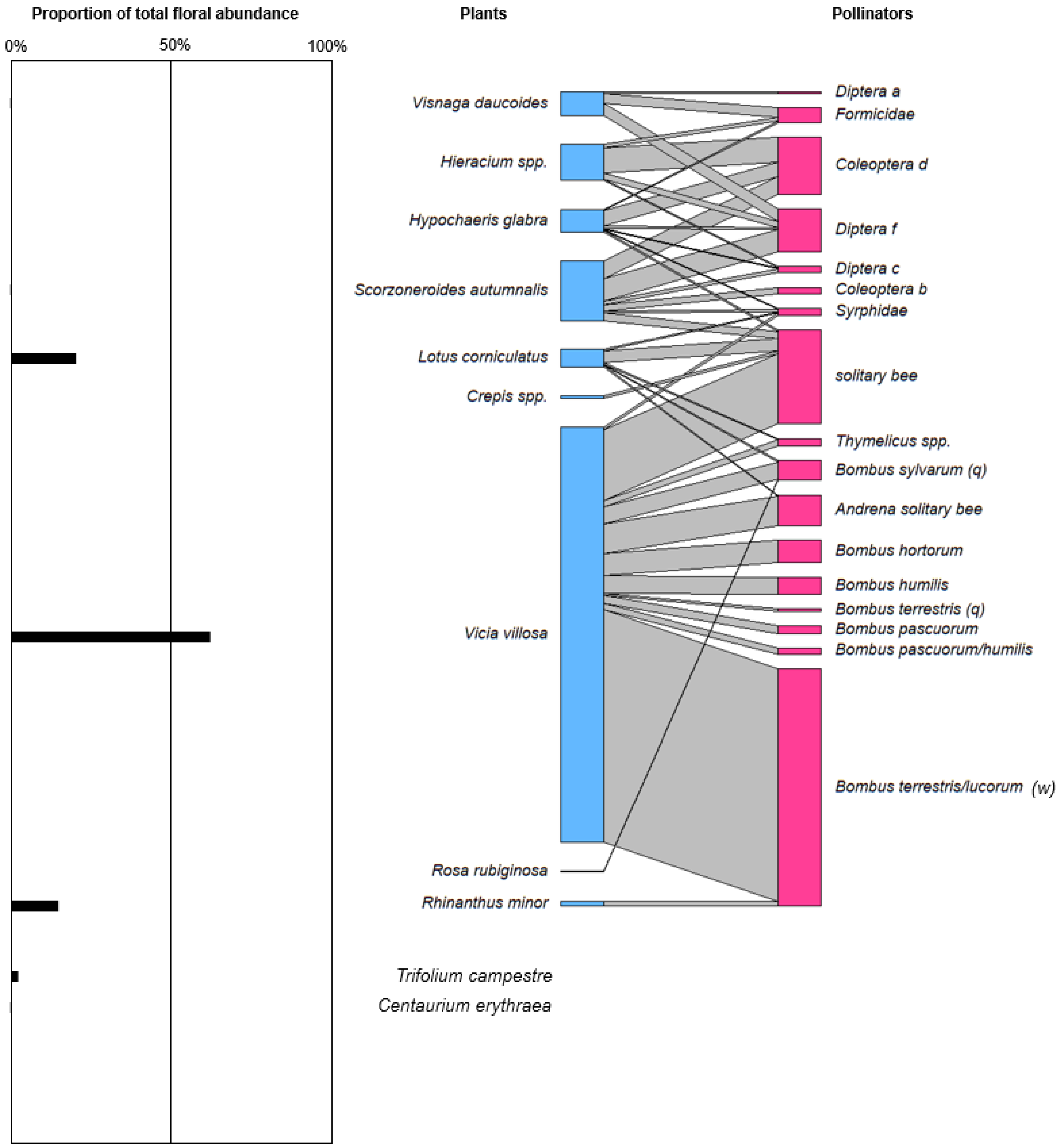
Proportion of total floral abundance
0%
50%
100%
Plants
Pollinators
Visnaga daucoides
Hieracium spp.
Hypochaeris glabra
Scorzoneroides autumnalis
Lotus corniculatus
Crepis spp.
Vicia villosa
Rosa rubiginosa
Rhinanthus minor
Trifolium campestre
Centaurium erythraea
Diptera a
Formicidae
Coleoptera d
Diptera f
Diptera c
Coleoptera b
Syrphidae
solitary bee
Thymelicus spp.
Bombus sylvarum (q)
Andrena solitary bee
Bombus hortorum
Bombus humilis
Bombus terrestris (q)
Bombus pascuorum
Bombus pascuorum/humilis
Bombus terrestris/lucorum (w)

**(d)**

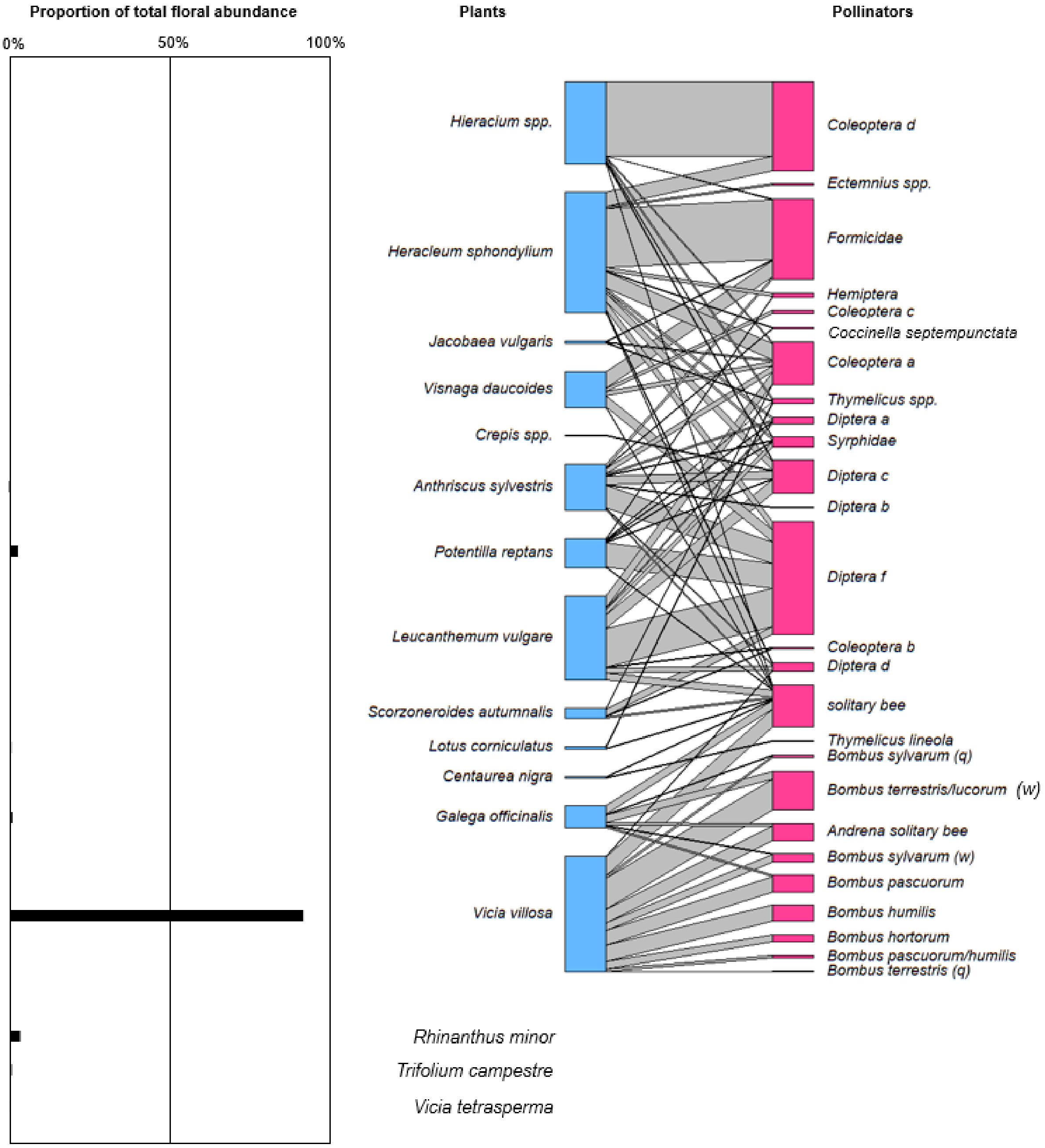


**Supplementary Fig. 1** (right) The interaction network between plants and pollinators recorded during transect walks at Coalhouse Fort, Essex, in May-June 2025 for each of the four meadows sampled: **(a)** meadow 2; **(b)** meadow 4; **(c)** meadow 5; **(d)** meadow 6; ‘q’ = bumblebee queens; ‘w’ = bumblebee workers; (left) the abundance of each plant species in each network: some plant species do not have abundance measures as they were not found in the quadrats along each transect (found in other places along each transect instead)
