## Supplementary Figure 2 for "Quantitative evidence for foraging preferences of shrill carder bumble bee (*Bombus sylvarum*) queens"

(a)

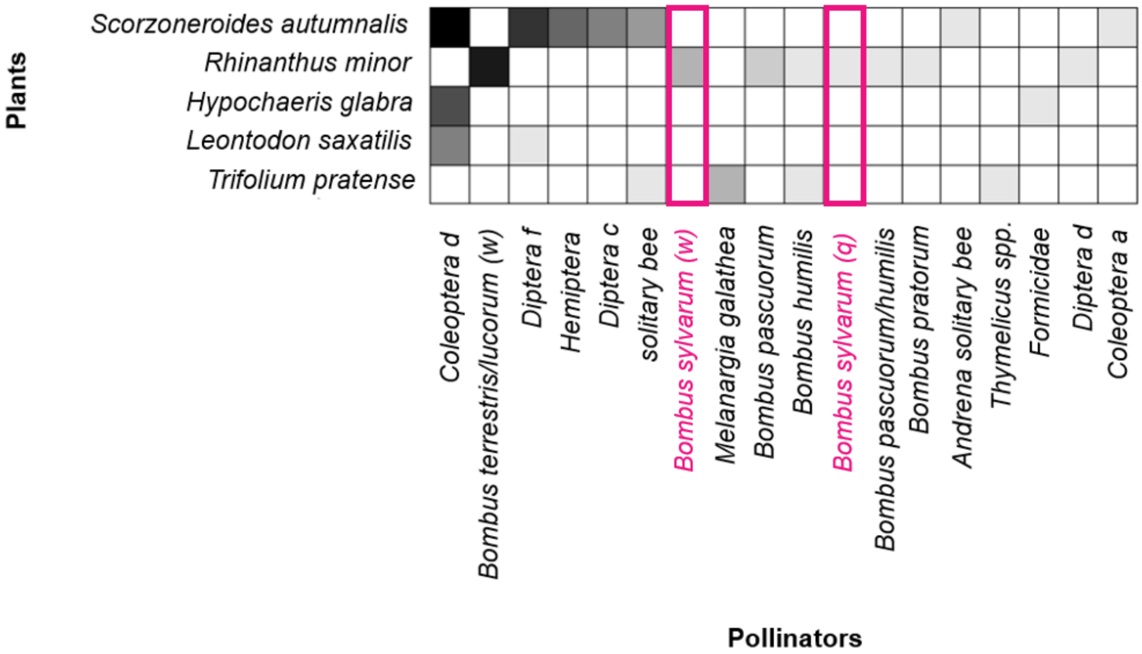
Plants
Scorzoneroides autumnalis
Rhinanthus minor
Hypochaeris glabra
Leontodon saxatilis
Trifolium pratense
Coleoptera d
Bombus terrestris/lucorum (w)
Diptera f
Hemiptera
Diptera c
solitary bee
Bombus sylvarum (w)
Melanargia galathea
Bombus pascuorum
Bombus humilis
Bombus sylvarum (q)
Bombus pascuorum/humilis
Bombus pratorum
Andrena solitary bee
Thymelicus spp.
Formicidae
Diptera d
Coleoptera a
Pollinators

(b)

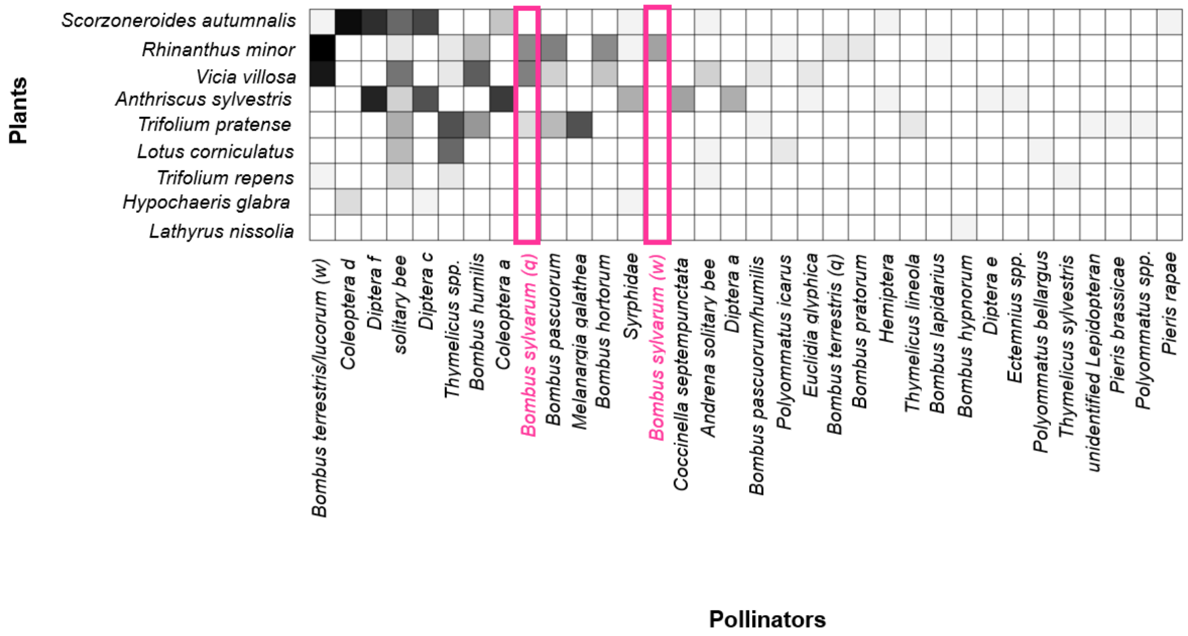
Plants
Scorzoneroides autumnalis
Rhinanthus minor
Vicia villosa
Anthriscus sylvestris
Trifolium pratense
Lotus corniculatus
Trifolium repens
Hypochaeris glabra
Lathyrus nissolia
Bombus terrestris/lucorum (w)
Coleoptera d
Diptera f
solitary bee
Diptera c
Thymelicus spp.
Bombus humilis
Coleoptera a
Bombus sylvarum (q)
Bombus pascuorum
Melanargia galathea
Bombus hortorum
Syrphidae
Bombus sylvarum (w)
Coccinella septempunctata
Andrena solitary bee
Diptera a
Bombus pascuorum/humilis
Polyommatus icarus
Euclidia glyphica
Bombus terrestris (q)
Bombus pratorum
Hemiptera
Thymelicus lineola
Bombus lapidarius
Bombus hypnorum
Diptera e
Ectemnius spp.
Polyommatus bellargus
Thymelicus sylvestris
unidentified Lepidopteran
Pieris brassicae
Polyommatus spp.
Pieris rapae
Pollinators

(c)

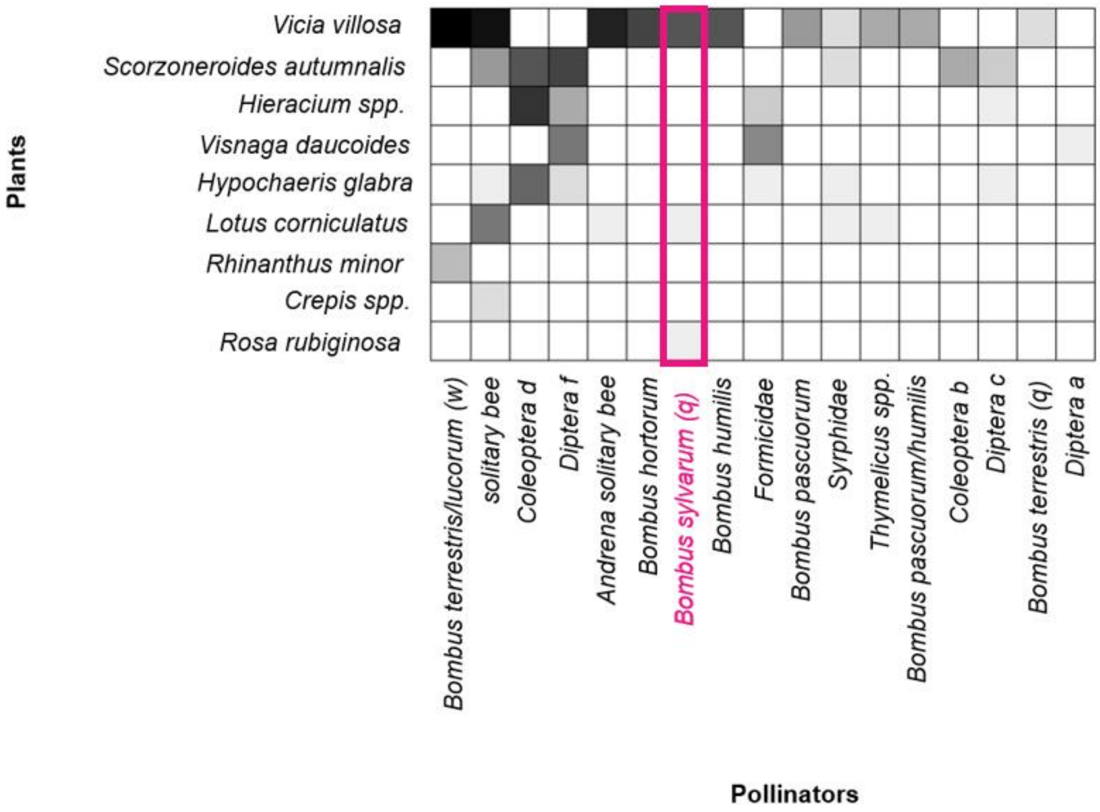
Plants
Vicia villosa
Scorzoneroides autumnalis
Hieracium spp.
Visnaga daucoides
Hypochaeris glabra
Lotus corniculatus
Rhinanthus minor
Crepis spp.
Rosa rubiginosa
Bombus terrestris/lucorum (w)
solitary bee
Coleoptera d
Diptera f
Andrena solitary bee
Bombus hortorum
Bombus sylvarum (q)
Bombus humilis
Formicidae
Bombus pascuorum
Syrphidae
Thymelicus spp.
Bombus pascuorum/humilis
Coleoptera b
Diptera c
Bombus terrestris (q)
Diptera a
Pollinators

**(d)**

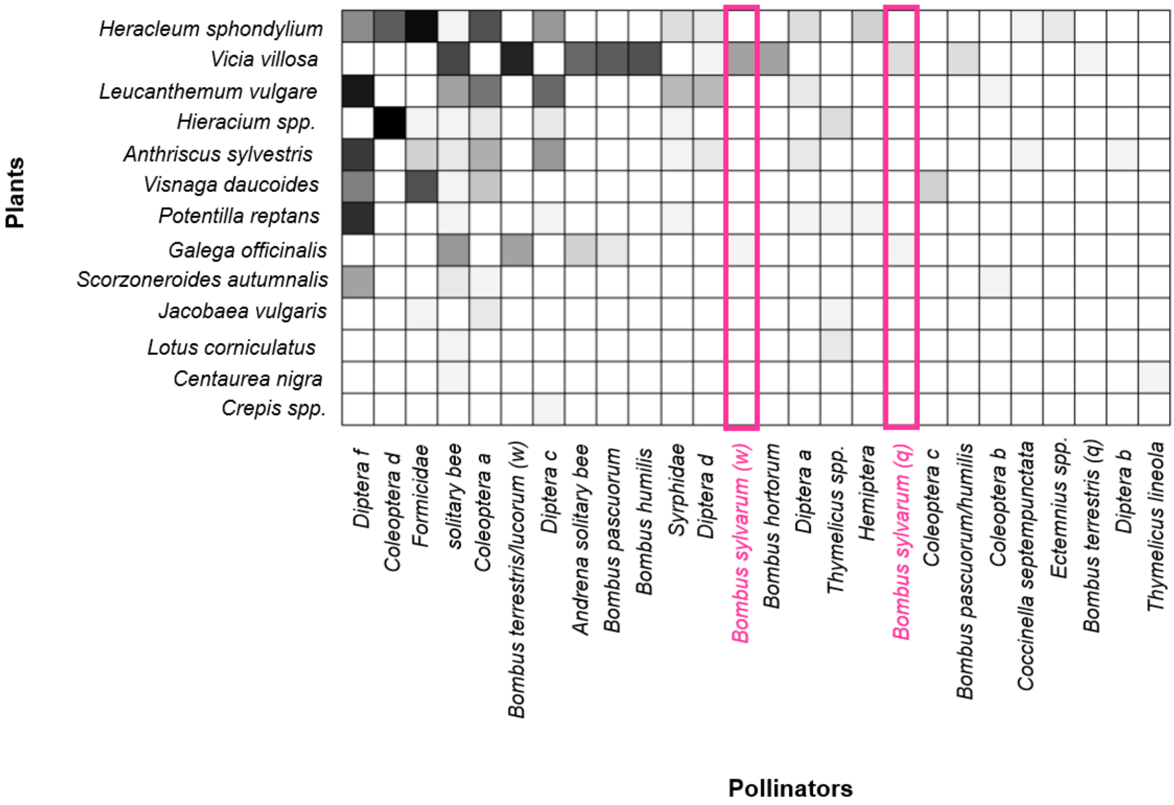


**Supplementary Fig. 2** Adjacency matrixes for the plants and pollinators seen interacting during the transect walks at Coalhouse Fort between May and June 2025 for each separate meadow sampled: **(a)** meadow 2; **(b)** meadow 4; **(c)** meadow 5, **(d)** meadow 6. The darker the cell, the greater the number of interactions between the two species. Empty cells are indicative of no interactions. *Bombus sylvarum* (queen ('q') and worker ('w')) interactions are highlighted
